# Developing best practices for genotyping-by-sequencing analysis in the construction of linkage maps

**DOI:** 10.1101/2022.11.24.517847

**Authors:** Cristiane Hayumi Taniguti, Lucas Mitsuo Taniguti, Rodrigo Rampazo Amadeu, Jeekin Lau, Gabriel de Siqueira Gesteira, Thiago de Paula Oliveira, Getulio Caixeta Ferreira, Guilherme da Silva Pereira, David Byrne, Marcelo Mollinari, Oscar Riera-Lizarazu, Antonio Augusto Franco Garcia

## Abstract

**Background:** Genotyping-by-Sequencing (GBS) provides affordable methods for genotyping hundreds of individuals using millions of markers. However, this challenges bioinformatic procedures that must overcome possible artifacts such as the bias generated by PCR duplicates and sequencing errors. Genotyping errors lead to data that deviate from what is expected from regular meiosis. This, in turn, leads to difficulties in grouping and ordering markers resulting in inflated and incorrect linkage maps. Therefore, genotyping errors can be easily detected by linkage map quality evaluations.

**Results:** We developed and used the `Reads2Map` workflow to build linkage maps with simulated and empirical GBS data of diploid outcrossing populations. The workflows run `GATK`, `Stacks`, `TASSEL`, and `Freebayes` for SNP calling and `updog`, `polyRAD`, and `SuperMASSA` for genotype calling, and `OneMap` and `GUSMap` to build linkage maps. Using simulated data, we observed which genotype call software fails in identifying common errors in GBS sequencing data and proposed specific filters to better handle them. We tested whether it is possible to overcome errors in a linkage map using genotype probabilities from each software or global error rates to estimate genetic distances with an updated version of `OneMap`. We also evaluated the impact of segregation distortion, contaminant samples, and haplotype-based multiallelic markers in the final linkage maps. Through our evaluations, we observed that some of the approaches produce different results depending on the dataset (dataset-dependent) and others produce consistent advantageous results among them (dataset-independent).

**Conclusions:** We set as default in the `Reads2Map` workflows the approaches that showed to be dataset-independent for GBS datasets according to our results. This reduces the number required of tests to identify optimal pipelines and parameters for other empirical datasets. Using `Reads2Map`, users can select the pipeline and parameters that best fit their data context. The `Reads2MapApp` shiny app provides a graphical representation of the results to facilitate their interpretation.

## 1 Introduction

Advances in sequencing technologies and the development of different genome-reduced representation library protocols result in millions of genetic markers from hundreds of samples in a single sequencing run (Glaubitz et al., 2014; Catchen et al., 2013; Anderson et al., 2018; Andrews et al., 2016). Increasing the number of markers and individuals genotyped can enhance the capacity of linkage maps to locate recombination events that occur, resulting in higher map resolution and better statistical power for the localization of QTL in further analysis. This large amount of data and genotyping errors common with genotyping-by-sequencing approaches (Bresadola et al., 2020) increases the need for computational resources and multiple bioinformatic tools.

Genotyping errors are frequent when high-throughput sequencing technology is applied to reduced representation libraries. There are a variety of protocols to create these types of libraries (Andrews et al., 2016), called Restriction-site Associated DNA sequencing (RADseq) or genotyping-by-sequencing (GBS) (Baird et al., 2008; Elshire et al., 2011). Generally, one or more restriction enzymes are used to digest the sample DNA. The resulting DNA fragments are filtered by size, connected to adaptors and barcodes, amplified by PCR, and sequenced. Consequently, most sequences obtained are PCR duplicates of the regions around the enzyme cut site. By relying on duplicates to increase sequencing depth, such methods introduce errors and a sequencing bias towards one of the alleles due to variabilities in the PCR amplification. These errors are hard to detect by bioinformatic tools (der Auwera and O’Connor, 2020; Rivera-Colón et al., 2020).

To overcome genotyping errors coming from GBS methods, genotype calling software model sequencing error, allelic bias, overdispersion, outlying observations, and the population Mendelian expected segregation (Gerard et al., 2018). Building a genetic map with genotypes obtained using these methods can be a powerful tool to validate their efficiency. Wrong decisions or inefficient methods in all steps before linkage map building can be identified in the resulting map as errors that dissociate the map properties from biological processes. For example, genotyping errors generate inflated map sizes that show an excessive number of recombination breakpoints during meiosis (a Hackett and Broadfoot, 2003). The first genetic map studies by Morgan and Sturtevant (Sturtevant, 1915) discovered that crossing-overs are unlikely to happen too close to each other, a phenomenon named interference. Later studies describing the meiotic molecular mechanisms confirmed the low expected number of recombination breaks in a single event (Smith and Nambiar, 2020).

Recently developed approaches to build linkage maps (Bilton et al., 2018; Mollinari and Garcia, 2019; Liao et al., 2021) were implemented in `OneMap` (Margarido et al., 2007) 3.0 package. They use quantitative genotype probability measurements rather than the traditional qualitative genotypic information from SNP and genotype calling methods to account for genotyping errors and provide higher-quality genetic maps. These probabilities can be applied in different ways: using the probability of each possible genotype (PL field in VCF format); using an error probability associated with the called genotype (GQ field in VCF format); or using a global error rate that will be applied to all genotypes. Nevertheless, even using these approaches, building a linkage map will succeed only if the upstream software can identify the errors and provide reliable genotypes or their probabilities.

The biallelic codominant nature of SNPs is another characteristic of high-throughput markers that can affect linkage map building of outcrossing species. Although biallelic markers can distinguish only two haplotypes, the mapping population of outcrossing diploid species inherits two haplotypes with combinations of four different parental haplotypes. With biallelic markers, the observed parental genotypes are limited to types *ab × ab, ab × aa*, and *aa × ab*. When one of the parents is homozygous (*ab × aa* and *aa × ab*), it is impossible to observe the crossing-over change for this uninformative parent. So this is taken as missing information (non-measurable crossing-overs) for linkage map building if only two-point information is considered. Therefore, building a linkage map with only biallelic markers requires a multi-point approach that uses loci information with both parents heterozygous (*ab×ab*) to estimate the recombination of loci where one parent is homozygous, and the recombination information is missing for closely linked loci. The multi-point approach applies likelihood computations involving several loci and has been successfully used since the seminal publication of Lander and Green (Lander and Green, 1987). The approach makes it possible to identify the four different parental haplotypes by phasing the biallelic information so that the SNPs can be used to identify all the allelic diversity.

Other approaches to overcome the low informativeness of biallelic markers involve combining adjacent biallelic markers in the same disequilibrium block (high LD) into a single multiallelic haplotype. These haplotype-based markers showed higher accuracy in association analysis than individual biallelic SNPs (Lorenz et al., 2010; Gawenda et al., 2015; N’Diaye et al., 2017; Sehgal and Dreisigacker, 2019; Abed and Belzile, 2019; Liu et al., 2008; Jiang et al., 2018). N’Diaye et al. (2017) and Jiang et al. (2018) pointed out several advantages of haplotype-based markers, including the higher capacity to identify epistatic interactions, the presence of more information to estimate identical-by-descent alleles and the reduction of the number of statistical tests to perform.

Despite the availability of many software for estimating genotype probabilities (Garrison and Marth, 2012; Catchen et al., 2013; McKenna et al., 2010; Garrison and Marth, 2012; Clark et al., 2019; Serang et al., 2012; Gerard et al., 2018) and haplotype-based multiallelic markers (Garrison and Marth, 2012; Patterson et al., 2015), there are no recommendations about which combination and choice of parameters are the best for building linkage maps. Therefore, this work evaluates the consequences of building maps by applying genotype probabilities and haplotype-based markers from different software and parameters. To achieve these, we implemented new features in `OneMap` (Margarido et al., 2007), a widely-used software for building maps. We also developed the `Reads2Map` workflow, a tool to help users select a bioinformatic pipeline that provides the best quality markers to build a linkage map for their dataset. Here, we performed tests with simulated and empirical data and were able to make recommendations to users to obtain better linkage maps in several situations, such as low and high-depth sequencing, with and without segregation distortion, contaminant samples, and multiallelic markers, and using different software to perform the SNP and genotype calling.

## 2 Material and Methods

We developed `Reads2Map` (RRID SCR_023593), a collection of bioinformatics workflows using Workflow Description Language (WDL) (Voss et al., 2017). It enables sequence alignment, SNP and genotype calling analysis, and linkage map construction. With `Reads2Map`, researchers have the flexibility to explore various software options and parameter combinations, enhancing the construction of linkage maps. The workflows are available in GitHub (Taniguti et al., 2022a) and in workflowhub.eu (Taniguti et al., 2022b,c).

The `EmpiricalReads2Map` workflow was designed to evaluate empirical (real) datasets; and the `SimulatedReads2Map` workflow, was to simulate and evaluate datasets (figure 1). Both are composed of sub-workflows that can be run independently, which increases usage flexibility. There are multiple options available for running WDL workflows. Some of them are Terra.bio platform (Terra.bio, 2020) and Cromwell Execution Engine (Voss et al., 2017).

**Figure 1:**
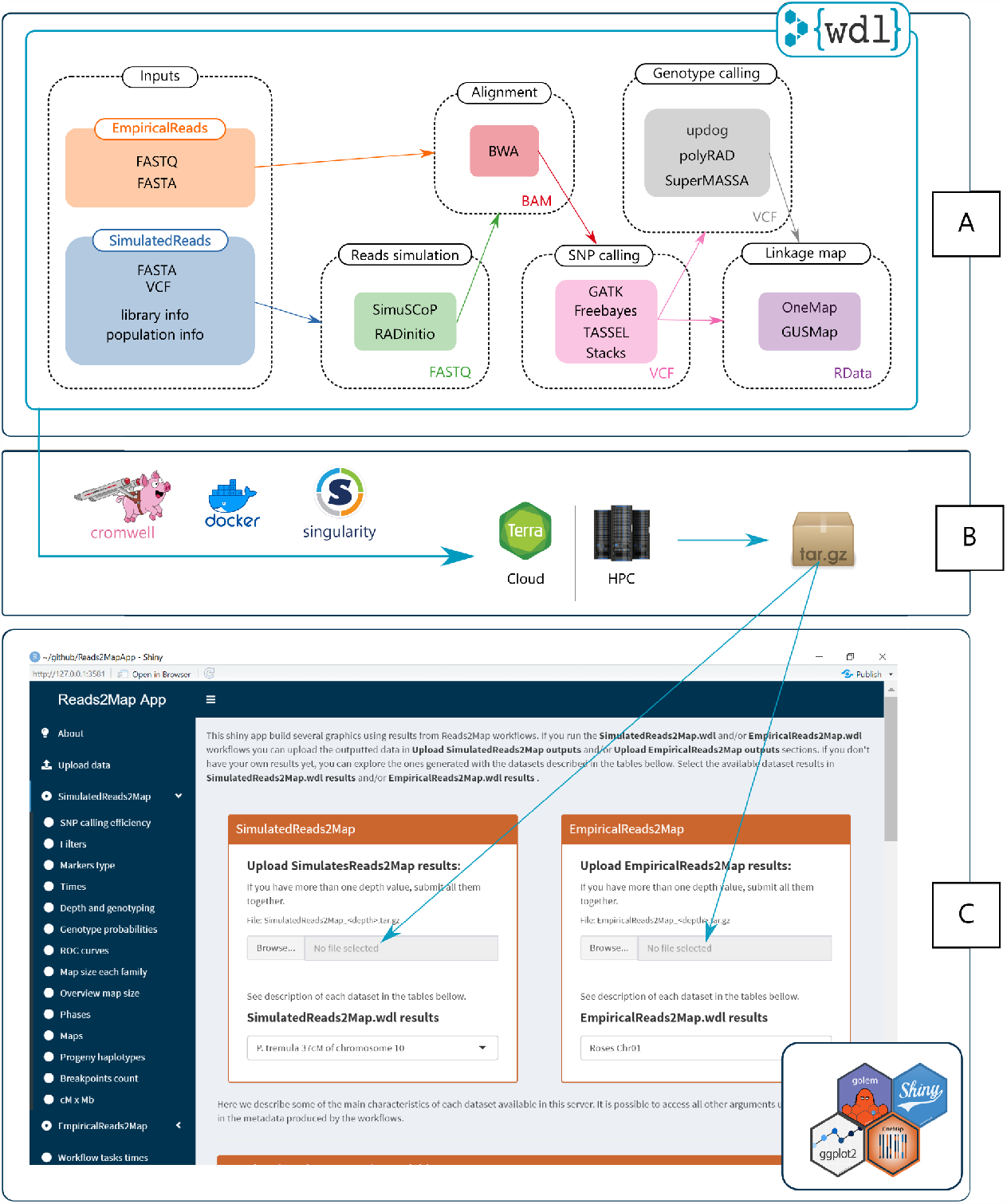
A: Tasks of the two main `Reads2Map` workflows: `EmpiricalReads2Map` and `SimulatedReads2Map`. B: Tools to run the workflows on the Cloud (Terra.bio, 2020) or in High-Performance Computing (HPC) environments. C: The `Reads2Map` shiny app has as input the outputs of the workflows. It builds several descriptive graphics to evaluate the best upstream software combination for linkage map construction.

Each WDL task in `Reads2Map` is related to a Docker (Merkel, 2014), or Singularity (Kurtzer et al., 2017) container. Some of the container’s images used in `Reads2Map` are available in open repositories and others were built using Dockerfiles stored in the `Reads2Map` repository and available in DockerHub. Check a list of all software and image versions used in Supplementary Table 1. We ran the analysis testing workflows on two high-performance computers (Texas A&M University HPRC, University of São Paulo Águia Cluster).

**Table 1:**
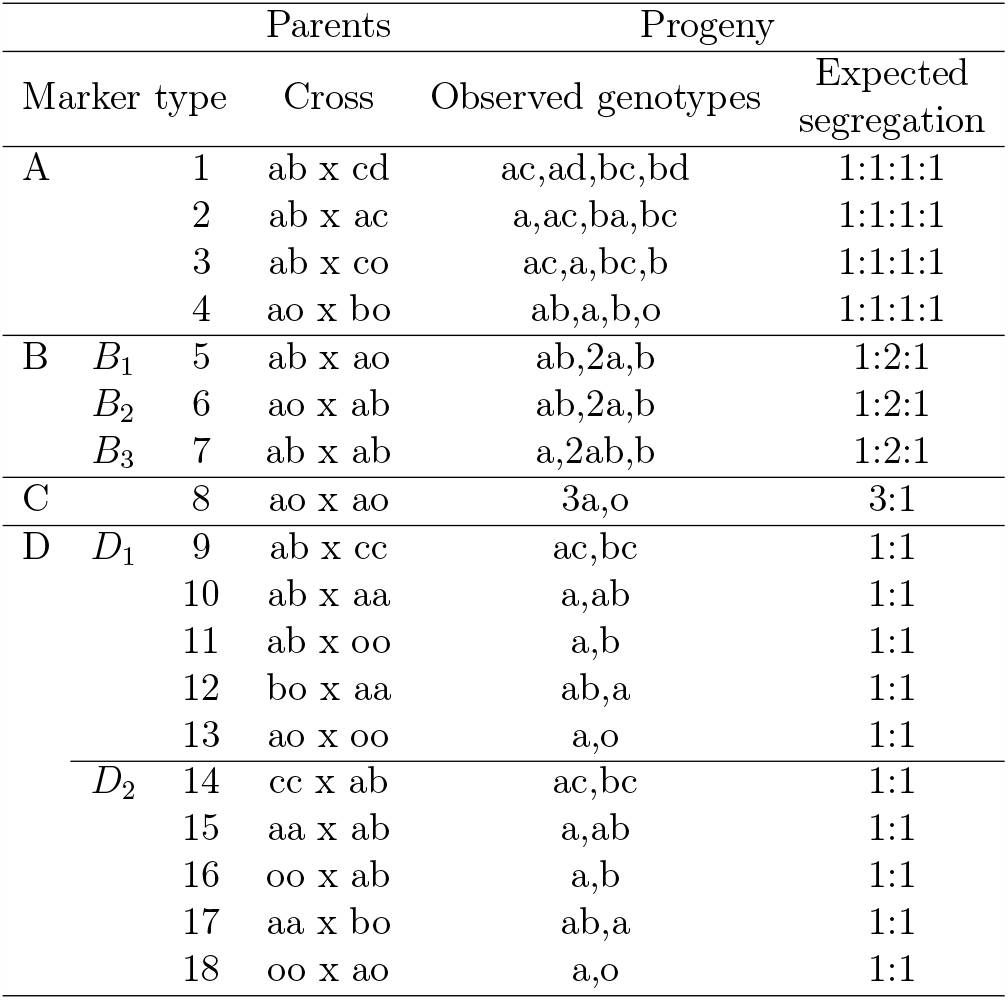
Marker types according to parental genotype combinations and progeny segregation. The letters “a”, “b”, “c” and “d” represent different alleles and the letter “o” represents null alleles. Adapted from (Wu et al., 2002).

For building linkage maps, we implemented updates in `OneMap` package version 3.0 and used this version in the workflows. We also developed the `Reads2MapTools` (Taniguti et al., 2022a) R package for support functions and `Reads2MapApp` shiny app (Taniguti et al., 2022b)), a visualization tool that receives as input the final workflow output and provides summary statistics about the resulting linkage maps, intermediary steps, and workflow performance.

### 2.1 SNP calling

The first step of the workflows is the SNP calling. To start with `GATK` (McKenna et al., 2010), `Stacks` (Catchen et al., 2013), and `Freebayes` (Garrison and Marth, 2012) approaches, the demultiplexed FASTQ sequences are first aligned to their respective reference genomes using `BWA-MEM` (Li, 2013). The workflow uses `samtools` (Li et al., 2009) to merge the alignment of replicates, keeping the libraries identification on the BAM header and filtering out reads with MAPQ < 10. After the alignment, BAM files for each sample are used as inputs for sub-workflows with `GATK`, `Stacks`, and `Freebayes` tasks. The `gatk_genotyping` subworkflow reproduces `GATK` joint genotyping via `HaplotypeCaller`, `GenomicsDBImport`, and `GenotypeGVCFs` tools and applies the suggested hard-filtering procedures (der Auwera and O’Connor, 2020). The `freebayes_genotyping` sub-workflow runs `Freebayes` parallelized by reference genome intervals. The `stacks_genotyping` sub-workflow includes the option to input the population file. If not included, all individuals are considered from the same population. It runs the `gstacks` and the `populations` plugins.

The `TASSEL` (Glaubitz et al., 2014) SNP caller is implemented in the `tassel_genotyping` sub-workflow. It first adds fake barcodes to the demultiplexed FASTQ sequences. After, it runs the plugins `GBSSeqToTagDBPlugin` and `TagExportToFastqPlugin`. The generated tags are aligned to the reference genome using `BWA-MEM` and the alignment files are input for the `SAMToGBSdbPlugin` plugin which produces a database. The database was processed by the `DiscoverySNPCallerPluginV2`, `SNPQualityProfilerPlugin`, and `ProductionSNPCallerPluginV2` plugins.

After obtaining the VCF file using one or more of the SNP calling methods, indel marker positions are left-aligned and normalized with `BCFtools` (Danecek et al., 2021).

### 2.2 Genotype calling

The VCF files with biallelic markers from `Freebayes`, `TASSEL`, `Stacks`, and `GATK` are the input for the genotype caller software `polyRAD` (Clark et al., 2019), `SuperMASSA` (Serang et al., 2012), and `updog` (Gerard et al., 2018). These three software are implemented in the sub-workflows `genotyping_empirical` and `genotyping_simulated`.

To use the `polyRAD` approach, the VCF files are imported using `VCF2RADdata` without applying any filters or considering phase information. The `polyRAD` model is run with `PipelineMapping2Parents` default arguments which assume an *F*_1_ bi-parental population. The function `Export_MAPpoly` is used to export the genotype probabilities. The `vcfR` package (Knaus and Grünwald, 2017) and custom R (function `polyRAD_genotype_vcf` in `Reads2MapTools` package) code is used to store outputted genotypes and their probabilities in a new VCF file. We also adapted `SuperMASSA` scripts to output the genotype probabilities information. The modified version is available in `Reads2MapTools` package. A wrapper function called `supermassa_genotype`, available in the package, can run the model in parallel and export the results to a new VCF file. The *F*_1_ `SuperMASSA` model is run with the parameter `naive_posterior_reporting_threshold` set to zero to not filter any genotype. The `updog` *F*_1_ model is used in parallel using the function `multidog` through the `Reads2MapTools` wrapper function `updog_genotype` which outputs the results in a new VCF file.

The software `GUSMap` performs the genotype calling and linkage map building with a single model. We use `VCFtoRA` function to convert the outputted VCF files from `GATK`, `TASSEL`, `Stacks`, and `Freebayes` approaches into `GUSMap` format. A pedigree of the population and a list of filters (MAF = 0.05, MISS=0.25, BIN=0, DETPH=0, and PVALUE=0.05) is provided to the `readRA` function. The function `makeFS` is used to create the full-sib population information. Functions `infer_OPGP_FS` and `rf_est_FS` are used to estimate the phase and recombination fraction given the genomic order of the markers. In some situations, the function `rf_est_FS` outputs infinite values of the recombination fraction. In these situations, our pipeline removes the respective marker and runs the function again. This workaround code can increase the time required to run `GUSMap`.

### 2.3 Updates in OneMap 3.0 for building linkage maps

`OneMap` is an open-source R package that has been serving the research community since its initial release in 2007. It offers a comprehensive suite of functions designed to facilitate marker filtering, grouping, ordering, and genetic distance estimation in both inbred and outbred populations. The genetic distance estimation is made using a Hidden Markov Model (HMM) multipoint approach. The forward-backward algorithm (Baum et al., 1970) is implemented to compute the HMM combined with the expectation-maximization algorithm (EM).

The `OneMap` latest version (3.0) is implemented in `Reads2Map` workflows. In this new version, we have introduced a new feature to enhance the flexibility of the HMM in scenarios where genotyping errors are expected in the dataset. This update includes the create_probs function and modifications to the HMM algorithm. With this option, users can provide `OneMap` with prior information regarding the reliability of each input genotype, thereby increasing the HMM’s adaptability. The create_probs function allows users to input three types of values: a global error value (global_error); an error probability for each inferred genotype (genotypes_error); or genotype probabilities for each possible genotype in individuals (genotypes_probs). This flexibility empowers users to tailor the analysis to their specific dataset characteristics and improve the accuracy of the results. This update is described in detail in Supplementary File 1.

The `OneMap` software previous to version 3.0 considered the HMM error probability as a single value of 10^*−*5^ for every genotype. In version 3.0, this value is kept as default to keep the code reproducible. However, it is noteworthy that this probability can be unreliable in several situations when the genotypes are more prone to errors, especially for new genotyping technology (e.g. GBS data).

`OneMap` 3.0 updates also include the possibility to parallelize the HMM using the approach described by Schiffthaler et al. (2017). It parallelizes the procedure into a maximum of four cores. We used this new `OneMap` feature to estimate the genetic distances. We also implemented new functions for linkage maps quality diagnostics such as interactive plots for recombination fraction matrices, progeny haplotype representation, and counts of the recombination breakpoints in progeny.

Despite using the parallelized HMM, the genetic distance estimations in `OneMap` can take time to run with a high number of markers, chromosomes, and tested combinations of software. Therefore, the `EmpiricalReads2Map` workflow runs the HMM in just a subset of markers which can be a single chromosome or a fragment of a chromosome. The alignment, the SNP, and genotype calling steps are performed with the entire dataset. After running the workflow and deciding the pipeline that provided the best results, the respective VCF output can be used to build the linkage map for all chromosomes in the R environment with `OneMap` functions.

The `OneMap` function `onemap_read_vcfR` is used to convert the VCFs to the `OneMap` R object format. The markers are filtered again by a maximum number of missing data of 25% because the VCF files include unexpected genotypes according to the segregation of a given locus (e.g. in a cross “AA x AB”, genotype “BB” cannot exist). `OneMap` makes this genotype call missing. Markers are also filtered if the segregation distortion is under a global significance level of 0.05 with Bonferroni correction and if they are redundant. Markers are ordered according to the reference genome position.

The `Reads2Map` workflows give flexibility to the user to define the probabilities to be used in the `OneMap` HMM for the estimation of the genetic distances. Users can provide more than one value to be tested as global errors (global_error input); can choose to use the upstream genotype caller error probability (genoprob_error input); and can provide global error values to be considered together with the software probabilities (genoprob_global_error input) according to the following: 1 *−* (1 *− global error*)*x*(1 *− software error probability*).

For `GATK`, `TASSEL`, `Stacks`, and `Freebayes` callers, the workflow uses in the HMM the Phred score genotype error (GQ FORMAT value) converted to probabilities. For the software `polyRAD`, `SuperMASSA`, and `updog` it uses 1*−output genotype probability* as a genotype error. For these last, the population’s structure (*F*_1_) is used as *a priori* information to increase the accuracy of the estimated genotypes.

The simulations do not consider interference in the recombination events. Therefore the Haldane map function was used to estimate the genetic distances in `SimulatedReads2Map`. Kosambi’s map function was applied to estimate the genetic distances in the `EmpiricalReads2Map`.

### 2.4 Read2Map Workflows App

The shiny app `Reads2MapApp` was built to display results from the workflow analysis. It includes graphics and statistics about SNP calling efficiency, the number of markers discarded by filtering steps, marker types, computer resources and time spent by each step of the workflow, allele depth by genotype, genotype probabilities, map size, map phases, recombination fraction matrix, progeny haplotypes, breakpoints count, and the correlation between linkage map and reference genome markers positions. `Reads2MapApp` is a modular R package using the `golem` framework (Guyader et al., 2022) that can be rendered and displayed locally or on a server. It can be installed from its GitHub repository and run with a single command (run_app). Once the `Reads2Map` output file is uploaded into the app, all graphics will be automatically generated.

### 2.5 Empirical datasets

We used the structure of `Reads2Map` to test the effects in the linkage map built using different combinations of software, and parameters in datasets with different characteristics. For our tests with empirical data, we used two datasets from previous works. They are GBS datasets from a bi-parental diploid *F*_1_ full-sib mapping populations of aspen (*Populus tremula L*.) (Zhigunov et al., 2017) (BioProject PRJNA395596), and rose (*Rosa spp*.) (Young et al., 2022). The aspen dataset comes from an intraspecific cross of two *Populus tremula* genotypes. The GBS libraries were built using *HindIII* and *NalIII* enzymes and sequenced as 150 base pair single-end reads on an Illumina HiSeq2500. Eight library replicates were built and sequenced for the parents and only one for each of the 116 F1 offspring. The dataset includes six samples erroneously sequenced as part of the progeny and later identified as contaminants. An average read depth of approximately 6x for progeny and 58x for parental samples was observed from the sequencing process. The *Populus trichocarpa* genome version 3.0 (Tuskan et al., 2006) was used as a reference for the sequence’s alignment. It is about 397 Mb in size. The diploid roses dataset comprises 138 individuals from the cross between a Texas A&M breeding line J06-20-14-3 (J14-3) and cultivar Papa Hemeray (PH). GBS libraries were built with *NgoMIV* enzyme and sequenced as a 113 base pair single-end read on a HiSeq2500. The parent J14-3 was repeated twice, and the PH sample three times. An average read depth of approximately 94x for progeny and 528x for parental samples was observed from the sequencing process. The *Rosa chinensis* v1.0 genome assembly (Saint-Oyant et al., 2018) was used as a reference genome to align the sequences. It is about 527 Mb in size.

The sequencing reads of the two empirical datasets were filtered using the `Stacks` plugin process_radtags (Catchen et al., 2013) to filter sequences by the presence of the restriction site and sequencing quality. The reads were discarded if the average quality score of 50% of its length was below the Phred score of 10 (or 90% probability of being correct). The software `cutadapt` (Martin, 2011) was used to remove adapters and filter by a minimum read length of 64 bp. The sequences were then evaluated in our `EmpiricalReads2Map` workflow.

### 2.6 Simulated GBS data

The first step of the `SimulatedReads2Map` workflow is to perform simulations of a mapping population, GBS libraries, and sequences. The simulation is based on a given reference genome chromosome sequence. If a reference linkage map and a VCF file are provided, the workflow simulates the marker genetic distances and parental genotype frequencies based on them. A cubic spline interpolation with the Hyman method (Hyman, 1983) is applied to simulate the centimorgan position for each marker’s physical position based on this same relation on the reference linkage map provided.

We based our simulation analysis on the first 37% of the chromosome 10 sequence of *Populus trichocarpa* version 3.0, which includes a sequence with 8.426 Mb from a total chromosome size of about 23 Mb. This sequence comprises 38 cM (21%) of the linkage group 10 built using the aspen empirical data (Zhigunov et al., 2017). Due to the computational resources needed to build such a high number of maps, we used only a subset of the data to finish the analysis in a reasonable time. Chromosome 10 was randomly chosen.

We simulated markers with different expected segregation patterns according to parental genotypes in each locus. Table 2 shows the notation for each possible marker type in an outcrossing diploid population. The `SimulatedReads2Map` workflow simulates parental haplotypes using the same proportion of marker types identified in the empirical VCF file. This approach overcomes the missing data present in the empirical dataset. The final VCF file used as a reference to the simulations contains 810 markers (126 B3.7, 263 D1.10, 278 D2.15, and 143 non-informative markers with both parents homozygous), which results from the aspen empirical data `GATK` SNP calling, filtered by a maximum of 25% of missing data and MAF of 5%.

**Table 2:**
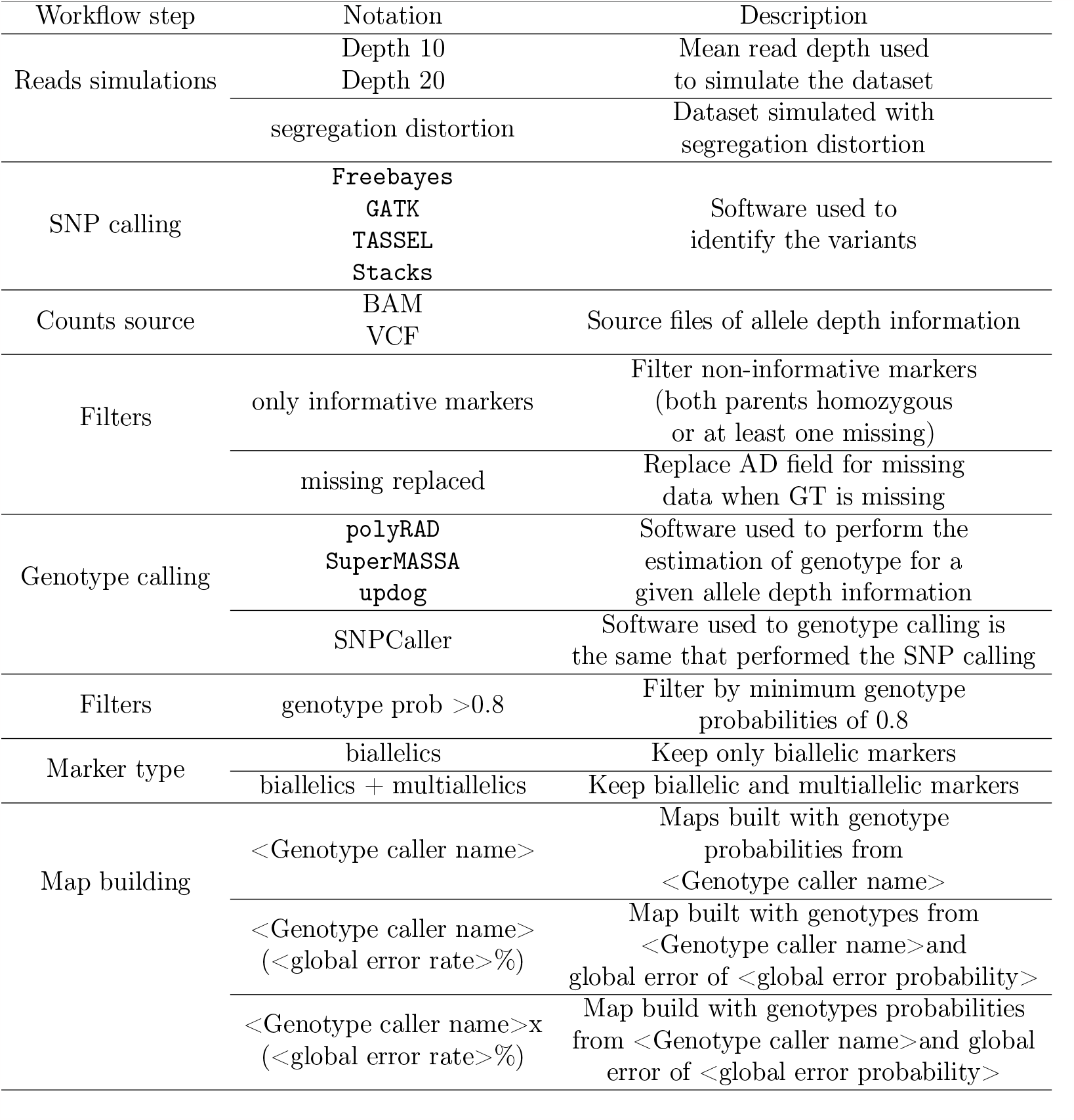
Notation used to refer to each evaluation scenario in empirical and simulated datasets.

`PedigreeSim` v2.1 software (Voorrips and Maliepaard, 2012) is implemented in the workflow to simulate the meiosis events and generate an *F*_1_ progeny based on the provided genetic map and simulated parental haplotypes. We did not consider interference in meiotic events (Haldane (1919) mapping function). `PedigreeSim` output files were converted to VCF files using `Reads2MapTools` R package function `pedsim2vcf`.

While converting the files, the `pedsim2vcf` function can also simulate segregation distortion by applying a selection strength. For that, a high number of individuals in the progeny have to be simulated with the `PedigreeSim` software and one or more loci to be under a given selection intensity. In our study, we targeted a final population size of 200 individuals. For that, we simulated 50 *×* 200 individuals and applied a selection intensity of 50% in the 30th marker, eliminating 50% of the genotypes containing one of the alleles. Then, 200 individuals of the resulting population are randomly selected to compose the mapping population. We used this feature to compare software performance in segregation distortion.

The VCF file output by pedsim2vcf and the reference genome file are inputs for the `RADinitio` (Rivera-Colón et al., 2020) software. `RADinitio` adds the VCF polymorphisms in the reference genome sequence and simulates the GBS sequences. It uses the inherited efficiency model (Best et al., 2015) to simulate a PCR-amplified pool of molecules. The model includes the heterogeneity of the PCR amplification and the polymerase substitution errors. Next, `RADinitio` applies the user-defined ratio between DNA original molecules to be sequenced and PCR duplicates to create a distribution that will define the number of times the pool of loci is sampled, the number of duplicate molecules that are generated from a RAD locus template, and the distribution of PCR errors in the resulting reads. We defined the default parameter with a proportion of 4:1. Besides the PCR errors inserted during the pool sampling, the software also includes a commonly observed error pattern, where the 3’ end of the read accumulates more errors than the 5’ (Glenn, 2011). We tested different values of PCR cycles (5, 9, and 14) and mean depth (5, 10, and 20) to simulate the FASTA files. We set the other `RADinitio` simulation parameters to obtain 150 bases of read length, sequence size of 350 (parameter “–insert-mean”), and restriction enzymes *HindIII* and *NalIII*. The mean read depth parameter for the parental samples was eight times higher than the progeny. The combination of `RADinitio` parameters that produced results closer to those observed in empirical data was selected to perform simulations with and without segregation distortion, five repetitions (five families), and two average sequencing depths (10 and 20) and 5 PCR cycles.

`RADinitio` does not output the sequence quality scores, so we converted the FASTA file format to FASTQ format, including a Phred score of 40 for every base simulated using `seqtk` (Li, 2020) software. After obtaining the FASTQ files, the `SimulatedReads2Map` workflow followed the same tasks as the `EmpiricalReads2Map`, with alignment, SNP and genotype calling, and linkage map build. The `SimulatedReads2Map` workflow makes comparisons between real and estimated results within each step. The comparisons made during the workflow can be visualized in the shiny app `Reads2MapApp`.

### 2.7 Tested scenarios

We ran all implemented software for SNP calling and genotype calling (`GATK`, `Freebayes`, `TASSEL`, `Stacks`, `updog`, `SuperMASSA`, and `polyRAD`) on the empirical and simulated datasets. In addition, we explored the substitution of VCF allele counts with counts from the alignment (BAM) files to mitigate potential biases introduced by SNP caller software when analyzing low-coverage sequence data. `GATK` inserts the bias when reads are filtered in the local reassembly step to avoid sequencing errors (Ros-Freixedes et al., 2017). `BCFtools` is used to find the read depths information for each allele in BAM files and update the allele depths information in the AD (allele depth) field of the VCF file. For the Aspen dataset, we also executed the workflows for every scenario in the presence of the contaminant samples.

The markers identified by the SNP callers (`GATK`, `TASSEL`, `Stacks`, `Freebayes`) were filtered by minor allele frequency (MAF) of 5% and maximum missing data allowed of 25% before proceeding to the genotype callers (`updog`, `polyRAD`, and `SuperMASSA`). At this step, we also tested two other filters. One of them was removing non-informative markers from the VCF file. We considered non-informative markers homozygous in both parents or if at least one of the parental genotypes was missing. The second filter was to replace the allele depth (AD) field in the VCF file format by missing data when the genotype is missing. This avoids that `updog`, `polyRAD`, and `SuperMASSA` use the allele depth when `GATK` filtered out the genotype due to bad quality.

After the genotype call, we reduce the analysis to a subset of markers (the first 8.426 Mb or 37%) of *Populus trichocarpa* chromosome 10 and the first 25 Mb (37%) of *Rosa chinensis* chromosome 1 reference genomes. This made it possible to build maps for all tests in a feasible time. The markers were filtered by the maximum missing data allowed of 25%, redundancy, and segregation distortion. In addition, we tested filtering the genotypes by a minimum genotype probability of 0.8.

We tested the consequences of building maps applying different genotype probabilities in the `OneMap` 3.0 HMM coming from seven different genotype caller software: `GATK`, `Freebayes`, `TASSEL`, `Stacks`, `polyRAD` (Clark et al., 2019), `SuperMASSA` (Serang et al., 2012) and `updog` (Gerard et al., 2018); a global error rate of 0.01, 0.05, 0.1, and the OneMap 2.0 default value of 10^*−*5^. We also tested the combination of the two distributions. We compared `OneMap` 3.0 capacity of estimating accurate genetic distances with the `GUSMap` package (Bilton et al., 2018) estimations since it also uses an HMM to account for errors present in sequencing data.

We also tested the consequences of the presence and absence of the and `Stacks` haplotypebased multiallelic markers in the linkage map. To test the influence of the presence of the multiallelic markers in the ordering procedure, we built a map for the entire chromosomes 1 and 10 from the roses and aspen datasets, respectively, using the selected pipeline. We ordered the markers using MDSMap (Preedy and Hackett, 2016) (wrapper function implemented in `OneMap` 3.0) ordering algorithm with and without multiallelic markers.

In the testing of scenarios in which we considered multiallelic markers, the VCFs containing them are merged into the VCF files from `polyRAD`, `SuperMASSA`, and `updog`. The merged VCF is the input for linkage map building in `OneMap` version 3.0.

Table 2 shows an overview of the notations used to refer to each evaluated scenario.

### 2.8 Performance comparison

We conducted performance comparisons of each tested dataset and scenario based on the built linkage map quality. To consider good quality we evaluate the following linkage map characteristics:

- Marker type: In outcrossing populations, it is important to have markers that have recombination information for both parents. We avoided approaches that provide only ab x aa (D1.10) or aa x ab (D2.15) in a single chromosome. The `Reads2MapApp` “Marker type” section describes the amount of each marker type in the linkage maps built by `Reads2Map` workflows.
- Marker coverage: It refers to how equally distributed markers are in the genome. We avoided approaches that do not detect markers in a large portion of the genomic selected area. The graphics in `Reads2MapApp` section “cMxMb” section correlate the linkage map position with the genomic positions. This is an excellent tool to evaluate marker coverage.
- Marker density: It refers to how equally distributed markers are on the linkage map. We avoided big gaps (higher than about 10 cM) in the linkage maps. Some of the gaps observed in the maps are due to outlier markers (a single marker with gaps in both edges). Outlier markers can be removed manually in further steps. We search for approaches that provided fewer outlier markers, which would require less manipulation later. The linkage map draw and graphics about the genetic distances among markers present in the section “Map size” of `Reads2MapApp` are good tools to evaluate marker density.
- Marker order: The efficiency of ordering algorithms can be significantly influenced by the presence of marker types that provide recombination information for both parents. In the `Reads2Map` workflows, to ensure accurate comparisons and to be possible to distinguish if linkage map inflation is due to different orders or genotyping errors, we have standardized the marker order across the workflow comparisons. Therefore, the order of the markers is always based on the reference genome. This means that it is crucial to carefully select, for the workflows, tests chromosome regions in the datasets that do not exhibit inversions or translocations when compared to the reference genome.
- However, in order to assess the impact of highly informative haplotype-based multiallelic markers, we conduct separate experiments outside of the workflows. In these experiments, we exclude outlier markers and evaluate the efficiency of the MDS ordering algorithms with and without the inclusion of multiallelic markers. This allows us to investigate these markers’ influence on the algorithm’s performance. We evaluated the orders provided by the different ordering algorithms by computing the absolute value of Spearman’s rank correlation between orders.
- Marker quality: In cases where all markers are correctly ordered (following the standardization in `Reads2Map` comparisons), and there is sufficient coverage and density, an inflated size of the linkage map can be attributed to a high error rate in the genotypes. Our objective is to find an approach that minimizes this inflation and brings the linkage map size closer to the expected value (e.g., 38 cM in our tested subsets).
- To identify the causes of inflated maps, the linkage map draw and recombination fraction matrix heatmap generated by `Reads2MapApp` prove valuable. It enables us to distinguish whether the inflation is a result of outlier markers creating gaps or due to genotyping errors.
- Estimated haplotypes: Together with the linkage map, the `OneMap` HMM multipoint approach also estimates the parents and progeny haplotypes. In a scenario without contaminant samples, we expect a low (around 1 or 2) and equally distributed number of recombination breaks across all samples. In scenarios where there are contaminant samples, we expect that their haplotypes contain a high number of estimated breaks because wrong assumptions were made leading to the wrong estimated number for these samples. `Reads2MapApp` contains a section for visualizing the progeny haplotypes and also for counting the estimated number of recombination breaks.

## 3 Results and Discussion

We use the structure of the `Reads2Map` workflows, the simulated, and the empirical datasets to test each software and some different parameters and markers filters. Our goal was to identify the approach that provides the best quality linkage map.

We have categorized the approaches used in our analysis into two groups: dataset-independent and dataset-dependent. The dataset-independent approaches consistently produce reliable results across all datasets, while the dataset-dependent approaches exhibit varying efficiency depending on the dataset characteristics. To streamline the user experience, we have selected the dataset-independent approaches that improve linkage map quality as the default options in the `Reads2Map` workflows (table 3). This simplifies the process for users by reducing the number of tests required, as these default approaches consistently yield favorable results across different datasets.

**Table 3:**
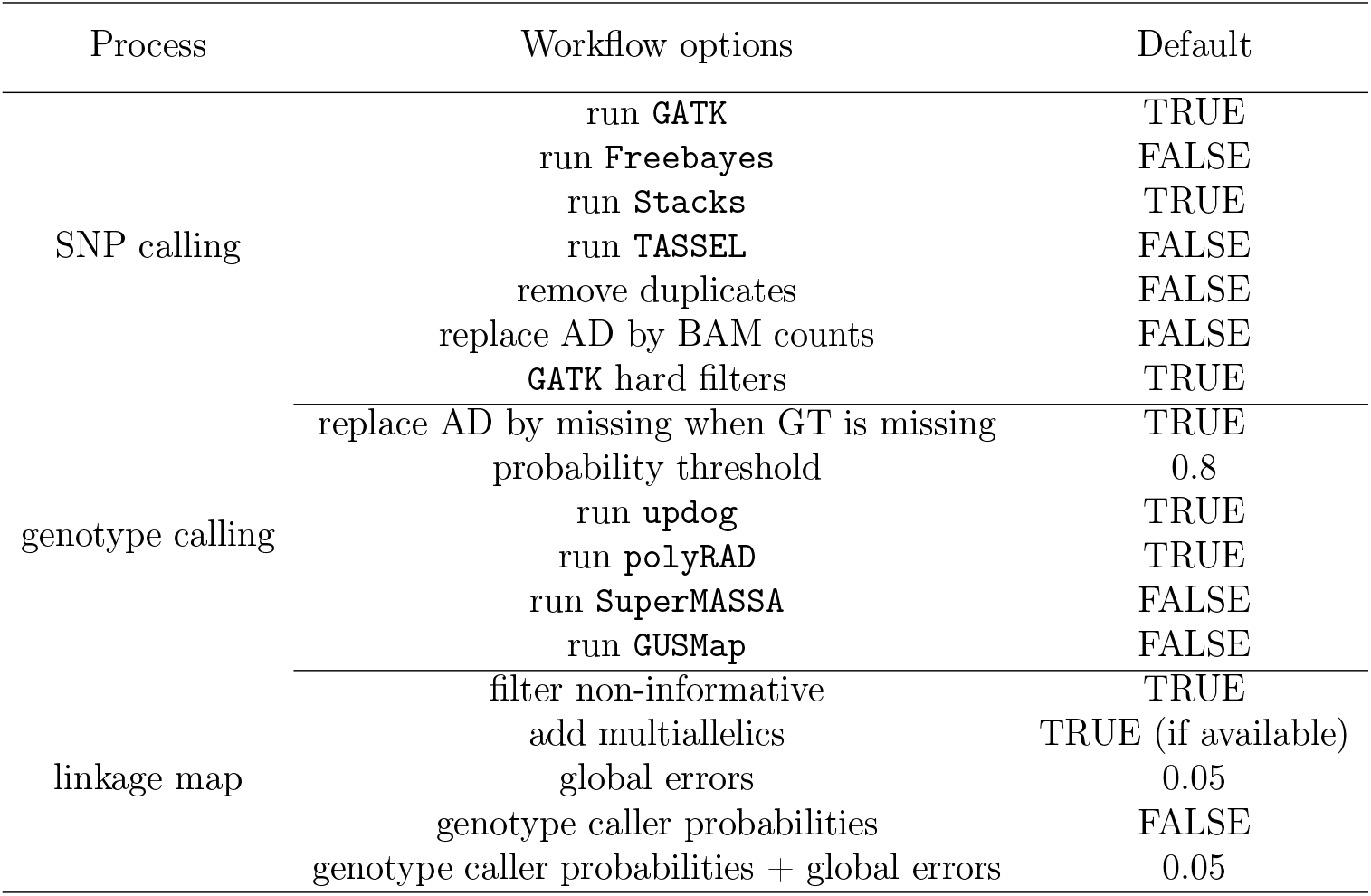
Reads2Map workflows default option set based on tests with empirical em simulated data.

We focused our tests and set the default options based on *F*_1_ diploid populations and GBS markers. However, because the `Reads2Map` workflow is modularized, the `EmpiricalSNPCalling` sub-workflow can be used separately and applied to other population structures, ploidy, and sequencing libraries. In the case of working with sequencing libraries other than RADseq, such as Whole-Genome-Sequencing (WGS) or Exome sequencing, it is important to set the option “remove duplicates” to TRUE. The PCR duplicates in RADseq data constitute the majority of the data and they are included in the allele count while calling the genotypes, but in other types of libraries, they are considered artifacts and are removed to avoid errors (DePristo et al., 2011).

The genotype call and linkage map building in the `EmpiricalMap` sub-workflow have the *F*_1_ population structure as an assumption. In this current version, they can be applied to another type of sequencing library but not to another type of population structure. For these steps, it is just important that the VCF file format is standardized and can be processed by BCFtools. They do not need to be necessarily from the SNP call software implemented. They can be also a combination of VCFs from different software such as the common markers between the implemented SNP call software results (“intersect” in Figure 2).

**Figure 2:**
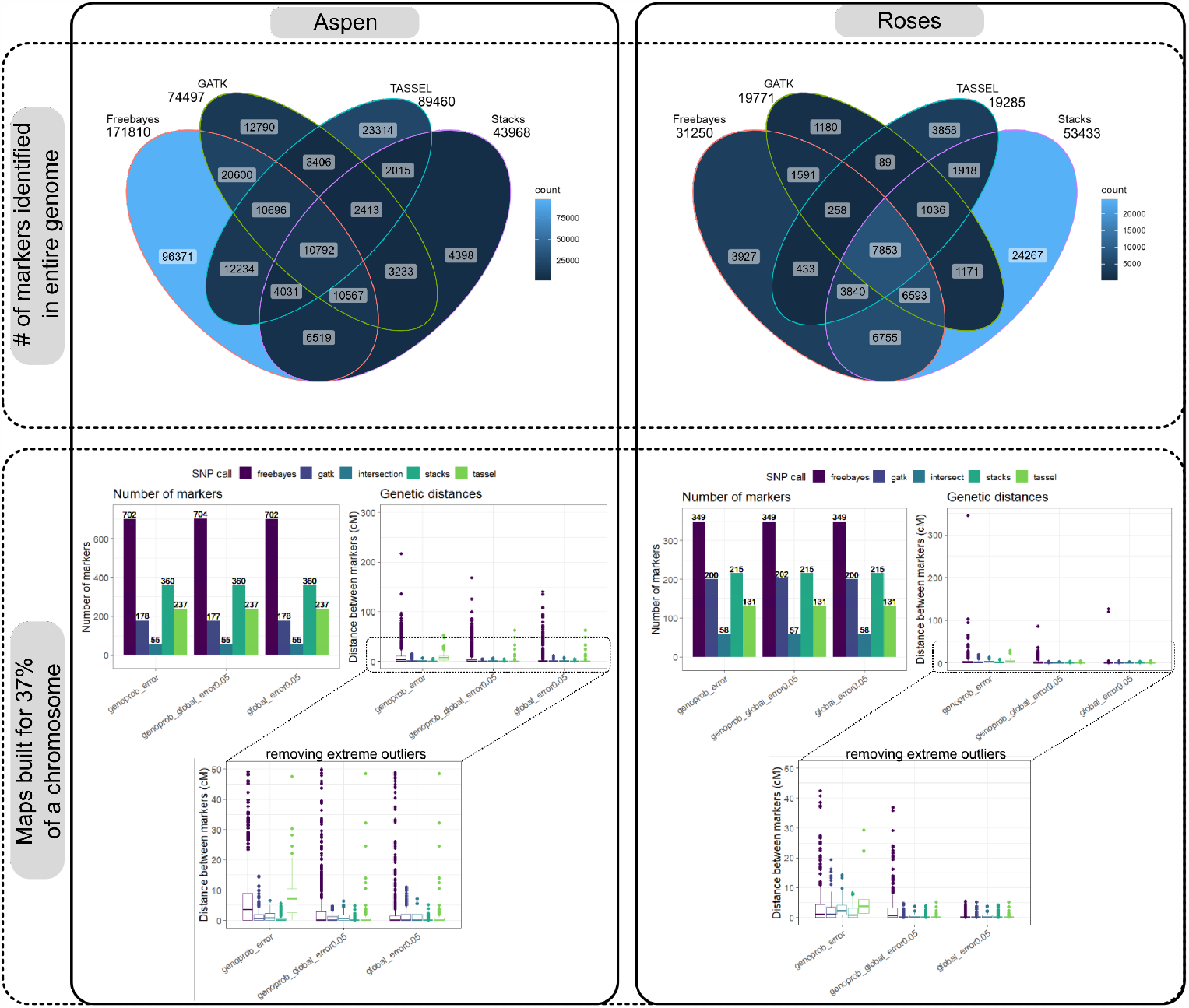
The top two figures show the number of markers identified by each SNP call software (number above each software name) and Venn diagrams showing the number of markers with common positions among all software results for the Aspen and Roses complete datasets. The markers were previously filtered by maximum missing data of 25% and MAF of 5%/. The compatibility of positions among markers from different software was only possible after using “BCFtools norm” to left-align the indels positions. The bottom two figures show the number of markers (bar plot) and distances between markers (boxplot) after building the linkage maps for a subset of 37% of chromosome 10 in the Aspen dataset and 1 in the Roses dataset with the markers from `Freebayes`, `GATK`, `TASSEL`, and `Stacks`. It was considered in the `OneMap` HMM the genotypes and a global error of 5% (global_error0.05); genotypes probabilities (genoprob_error); and the combination of genotype probabilities and a global error of 5% (genoprob_global_error0.0.5). These figures can be generated for user-defined empirical datasets in the `Reads2MapApp` sections “SNP calling efficiency” and “Map size” after running the EmpiricalMaps workflow.

We had to perform extra manipulations in `TASSEL` VCF output to be able to run the downstream analysis because they presented missing header information. Also, processing `Freebayes` showed to consume an unexpectedly high amount of RAM memory in some situations, which made it impossible to automatize the amount of memory required from the HPC and Cloud by the workflow task.

The number of markers identified by each software is related to the species, library preparation, and sequencing aspects such as genome size, restriction enzyme used, and sequencing depth. In figure 2, we can observe that more markers were identified in the Aspen dataset compared to the Roses due to the higher frequency of enzyme cut sites. There is no consistency between the two datasets about which of the software identifies the higher number of markers.

After all the filtering steps and linkage map building, it is consistent that `Freebayes` keeps more markers. However, the resulting maps built with `Freebayes` markers, genotypes, and genotypes probabilities presented higher genetic distances inflation compared to the other approaches. Using `TASSEL` software markers also resulted in higher inflation in Aspen dataset maps which have lower sequencing depth (∼ 6x) compared to the Roses (∼ 94). The other approaches also presented outlier markers that inflate the total map size, but, because they are individual markers, they can be easily removed in further steps. The maps built with only common markers among all four software (intersection in figure 2) contained fewer markers and had markers distances similar to `GATK` and `Stacks` results.

Evaluating the results of our simulations for `GATK`, we identified a format characteristic of VCFs from this software that leads to genotyping errors in estimations by `updog`, `polyRAD`, and `SuperMASSA`. In such cases, the genotype is considered missing in the `GATK` output VCF GT format field, while the total read depth is always reported in the reference allele field of the AD format field (e.g., Estimated = GT:AD ./.;22,0 | True = GT:AD 1/1;0,22).

We present examples of the consequences of this format in genotypes called by `updog`, `polyRAD`, and `SuperMASSA` in figures 3 and 4. In figure 3 A, allele dropouts are observed in the genotype of parent P2 and some of the progeny individuals. In empirical data, allele dropout can occur due to various reasons, such as polymorphisms in the cut site or the non-amplification of one allele during the PCR step (Rivera-Colón et al., 2020). Our simulations also consider allele dropout, but in the observed scenario, the source of allele dropout is due to the format characteristic of the `GATK` VCF file.

**Figure 3:**
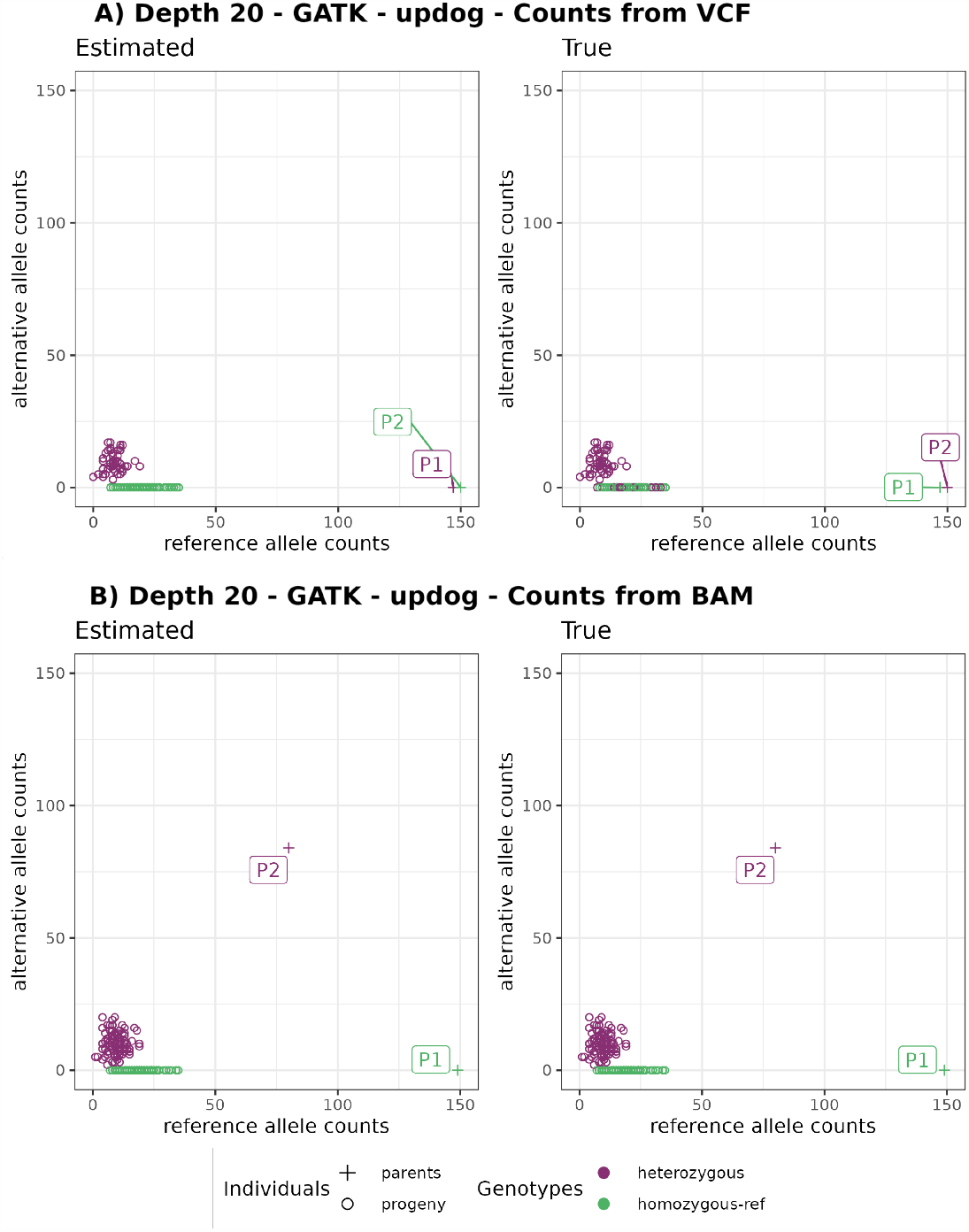
Example of error (Est: homozygous | True: heterozygous and Est: heterozygous | True: homozygous) in parental genotypes leading to a wrong marker type (Est: D1.10 | True: D2.15). Estimated reference (x-axis) and alternative (y-axis) allele count. Graphics on the left have colors according to estimated genotypes, and on the right to the true genotypes. A) show counts from `GATK` VCF file and B) from BAM file. In the VCF file outputted by `GATK` the P1 genotype is missing (GT ./.) because the reads did not pass the quality filters, but it reports the counts in the reference AD field (149,0). The `updog` software use progeny segregation (1:1) to estimate the parents, but it makes a mistake identifying which one is heterozygous. Using counts from BAM file (B) fix this issue despite losing the `GATK` quality filters that can be important in other situations.

**Figure 4:**
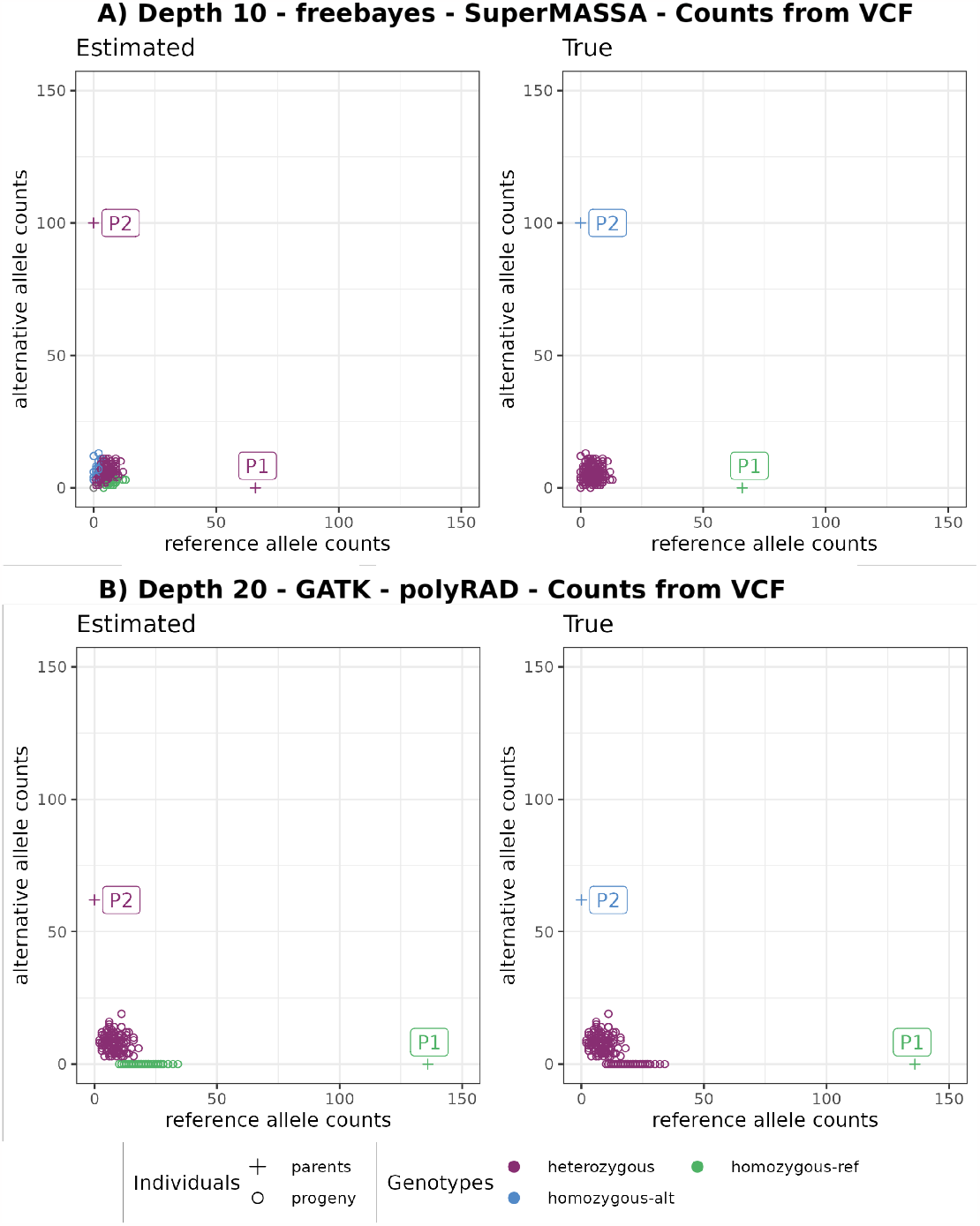
Example of error (Est: homozygous | True: heterozygous) in progeny genotypes leading to wrong marker types in A) Est: B3.7 | True: non-informative and in B) Est: D1.10 | True: non-informative. Graphics on the left have colors according to estimated genotypes, and on the right to the true genotypes.

The occurrence of genotyping errors while using `GATK` VCF allele counts was previously observed by Ros-Freixedes et al. (2017), who suggested using counts from BAM alignment files to address the issue (Figure 3 B). However, when testing the usage of BAM allele counts, we lose the advantage of the robust filtering applied by the `GATK` pipeline to retain only high-quality read counts in its VCF allele depth field. To maintain the accuracy of the `GATK` allele depth while overcoming the common error observed when the genotype is missing, we replaced the VCF allele count (AD and DP fields) with zero when the genotype information is missing before utilizing it for genotyping with `polyRAD`, `SuperMASSA`, and `updog`. This more precise way of solving the issue was only possible due to our simulations studies once they provide a clear comparison between simulated (true) and estimated data which highlighted the sources of the genotyping errors.

We also observed situations in `updog`, `polyRAD` and `SuperMASSA` results where the parental genotypes are wrongly estimated because of the low quality of the progeny genotypes that distort the expected segregation. These genotype call software consider the expected segregation in their models therefore errors in the progeny leads to errors in the parents. Figure 4 shows examples where the marker would be considered non-informative for an outcrossing population, as both parents are homozygous. However, due to genotyping errors in the population, `SuperMASSA` and `polyRAD` incorrectly estimate the parents as heterozygous. To tackle this problem, we implement a filtering step to exclude non-informative markers before applying the genotype callers.

Solving these issues was particularly important because erroneous parent genotypes have a higher impact on linkage map quality than progeny genotype errors. `OneMap` 3.0 does not consider the parental genotype probabilities in its HMM multi-point approach. Thus, it is important to plan the sequencing experiment with high-quality parental genotypes because, if there are errors, they will not be corrected in downstream processing, and it will cause distortions in the resulting distances and haplotypes. To avoid map size inflation, erroneous parental genotypes must be removed before the linkage map analysis.

In general, the evaluations of `RADinitio` simulations profile shows that we can expect fewer markers and genotyping errors in the simulated compared to the empirical data (Supplementary Figure 7). A smaller number of markers should not reduce the built linkage map quality because the analysis was made in *F*_1_ populations, which have large disequilibrium blocks. However, the smaller number of genotyping errors overestimates the SNP and genotype calling software efficiency. This overestimation is commonly observed in simulation results once the data cannot capture all biases and errors in the empirical data. Thus, we used the simulations to understand specific software limitations and error sources but not ultimately define the best performance (Duncavage et al., 2022).

We observed the same or improved quality of linkage maps in the empirical datasets evaluations (Supplementary Figure 8) when we applied these two described filtering steps: removing non-informative data before genotype calling, and replacing allele counts with missing data when the genotype is missing in the `GATK` calls. After the genotype calling, we applied a threshold of 0.8 to filter low-quality genotypes, which also was beneficial in all scenarios. It is important to notice that these filters are applied before the segregation test filter, which reduces the number of tests and increases the permissibility of the threshold corrected by multiple tests (Bonferroni correction). Thus, the built map can have more markers in some scenarios even if more filters are applied.

The simulations were also useful to validate all code developed for the analysis and to measure the effects of segregation distortion. The results showed that the segregation distortion does not affect the frequency of correct estimated genotypes in most scenarios, despite affecting the reliability of the genotype probabilities provided by `updog`, `SuperMASSA`, and `polyRAD` (Supplementary Figures 9 and 10). This can be one of the reasons why using genotype probabilities in the HMM did not present consistent results across tested datasets. Despite we considered the HMM error rate dataset-dependent values, we identified that some of the possible values can be discarded. Using the `OneMap` default value of 10^*−*5^ global error rate produced bad-quality maps in all situations. The same happened while using all the genotype call software relative error. Using higher values of global error rate and genotypes from `GATK`, `Freebayes`, `TASSEL`, `Stacks`, `updog`, and `polyRAD`, or the combination of the genotype probability and a global error rate from software `GATK`, `updog`, `Stacks`, and `polyRAD` produced the most reliable linkage maps, with linkage map sizes closer to the expected.

As observed in figure 5, many of the approaches produced linkage maps with distances between all adjacent markers smaller than 10 cM. We chose the method that results in less inflated linkage maps and outlier markers even when applying the small values of the global error rate (0.01). Once the method was selected, we tried an intermediary global error rate (0.075) for the roses dataset values to adjust to the expected total size. We also checked the recombination fraction heatmap, the markers coverage, density, and the number of estimated recombination breakpoints in progeny through `Reads2MapApp` figures (see the app interface demonstration in Supplementary File 2).

**Figure 5:**
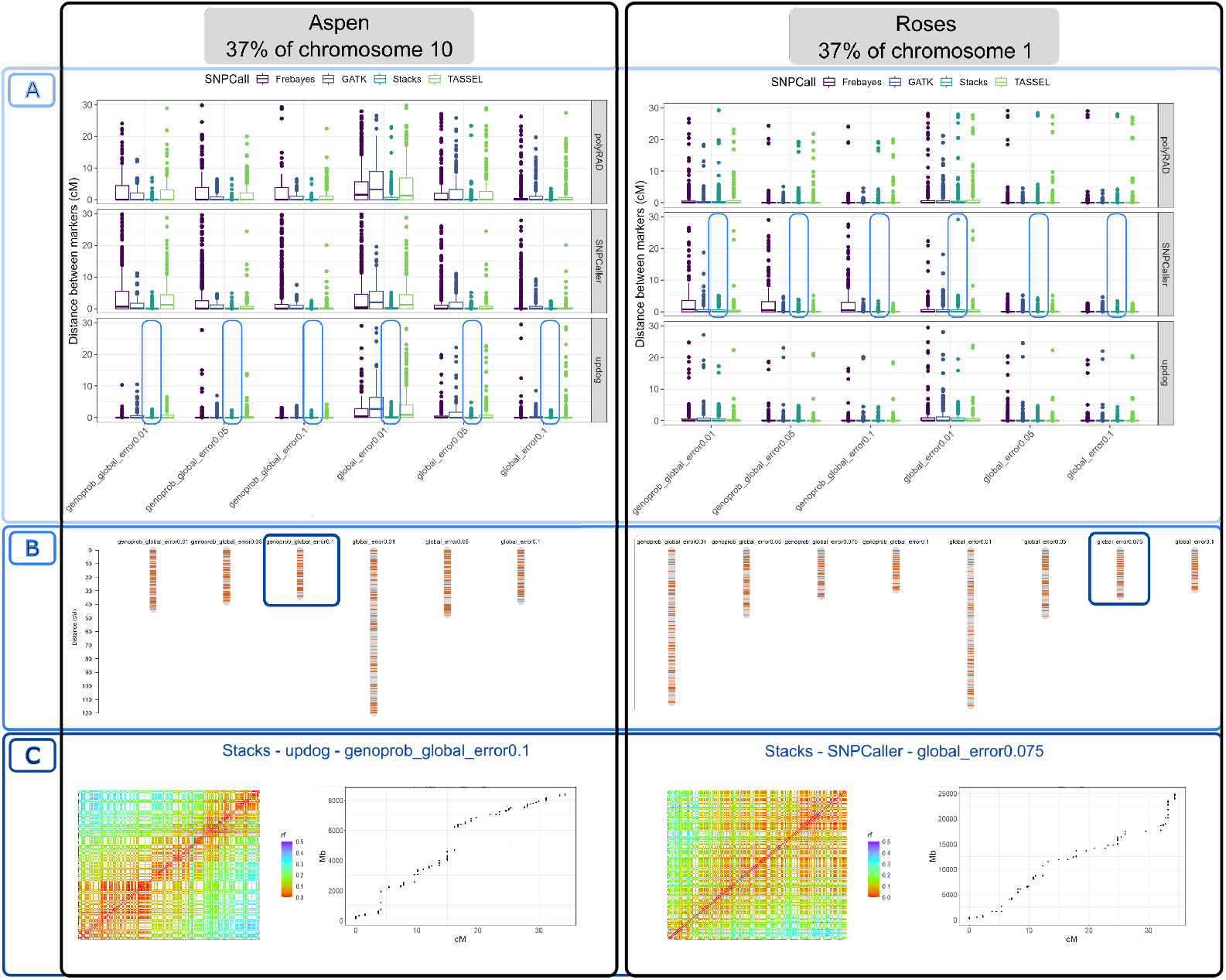
Process of selecting best pipeline: A) Comparing the effect of different error probabilities in the `OneMap` 3.0 HMM in the distances between adjacent markers; B) Comparing the effect of different error probabilities in the linkage maps total size built with a single SNP call software; C) Checking the recombination fraction (rf) heatmap and markers coverage in the genome using the selected pipeline. These figures were extracted from `Reads2MapApp`.

Before using the map size as a metric for map quality, we checked if a map with the expected size always means good quality. A map can have the expected size but a poor quality if the number of overestimated and underestimated recombination breakpoints in the progeny haplotypes is the same; in other words, if they cancel out. To test if this happens in our simulated dataset, we compared the Euclidean relation of estimated and true genetic distances with the total number of wrong (overestimated + underestimated) recombination breakpoints in the progeny haplotypes (Figure 6). For identifying a break as overestimated or underestimated, we do not consider the expected break position but the total breaks expected for the evaluated haplotype. For example, if one haplotype for a specific progeny was simulated with one break and estimated with zero, then we count it as one underestimated break.

**Figure 6:**
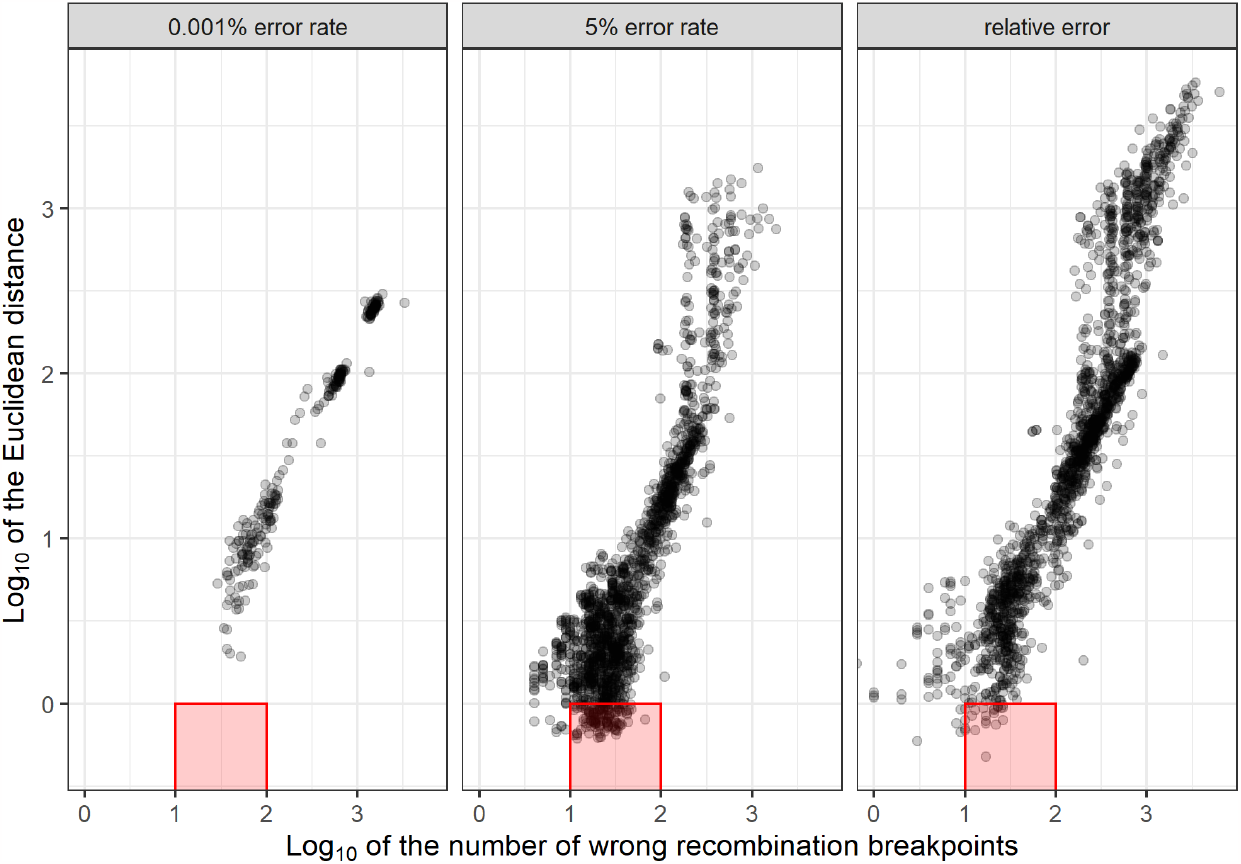
Relation between Euclidean distance (y-axis) and the number of recombination breakpoints (x-axis) in maps built with global error rates (0.001% and 5%), and with probabilities outputted by the genotype call software (relative error). Each dot represents a map built with simulated data based on the first 37% of aspen chromosome 10. The red squares highlight maps that do not present inflated size (1 or less Euclidean distance) but have from 10 to 100 wrong recombination breakpoints.

The comparison shows that overestimated breakpoints are generally more frequent than underestimated ones. We observe that when a map is inflated, it also has many wrong recombination breakpoints. However, in some cases, the map has the expected map size, but a high number of wrong haplotypes due to both overestimated and underestimated breaks. A high number of underestimated breaks can be observed in situations where the Euclidean distance is close to, or less than 1 (*log*_10_0) and the number of wrong recombination events is between 10 and 100 (*log*_10_1 and *log*_10_2). These situations are more frequent when a global error rate of 5% is used.

In the empirical data results, we observed maps with expected size and excess recombination breakpoints in just a few individuals in the progeny. This variation can be related to contaminant samples. The study of Zhigunov et al. (2017) identified six contaminants in the Aspen dataset. When we ran the workflows, including the contaminant samples, the maps built with `Freebayes` markers and `updog`, `SuperMASSA`, and `polyRAD` were smaller in size than without the contaminant (Supplementary Figure 11). This would (wrongly) suggest better quality if map size is the only metric used. Nevertheless, the maps presented higher differences in the number of recombination breakpoints among individuals when using the genotype probabilities relative to each genotype call software. Some contaminant samples presented more estimated recombination events than the rest of the progeny. Using higher values of global error reduces this difference and can mask the presence of contamination.

These results show that it is important to exclude contaminant samples before the linkage map building once the multi-point HMM approach tends to fix the genotypes according to the biological assumption that they are all *F*_1_ individuals. There are several methods available for identifying contaminant samples in previous steps. The ADMIXTURE (Alexander et al., 2009) software analysis as made by Zhigunov et al. (2017) is one possibility. Another is to calculate a marker-based relationship matrix using the R package AGHmatrix (Amadeu et al., 2016).

So far, all the evaluations we have discussed have focused exclusively on biallelic markers. We also evaluate the impact on the genetic distances when haplotype-based multiallelic markers are included. In most of the tested scenarios, incorporating these markers leads to map inflation. This is primarily due to the fact that inaccurately estimated multiallelic markers or genotyping errors associated with them can significantly affect the quality of the linkage map. The impact is particularly pronounced because multiallelic markers provide richer information, including recombination and phase information for both parents, compared to biallelic markers. However, the advantages of including the multiallelic markers appear in the marker ordering step.

Algorithms that use two-point recombination fractions estimations have issues ordering only biallelic markers because of the missing linkage information between markers D1 and D2 (homozygous x heterozygous or vice-versa). These markers can only be related to each other in the presence of more informative markers, such as B3.7 (heterozygous x heterozygous) or multiallelic states. Yet, having few B7.3 markers compared to D1 and D2 can still be an issue for linkage map building. In fact, this characteristic was the reason behind the initial development of separate maps for each parent in the first methods used for building genetic maps in such populations (Grattapaglia and Sederoff, 1994). These non-integrated genetic maps subsequently limited further analysis of multiallelic traits in terms QTL mapping (Gazaffi et al., 2014).

The markers ordering efficiency is not considered by `Reads2Map` workflows once it uses the genomic order to position the markers in the linkage maps. The reference genome is a required input by the workflows to standardize the positions of the markers across all tested methods. This avoids the confounding interpretation of bad-quality linkage maps due to wrong ordering and not genotyping errors.

To test the effect of multiallelic markers in the ordering, we built a linkage map for the entire chromosome 1 and 10 of the roses and aspen datasets, respectively, using the selected methods and adding the haplotype-based multiallelic markers provided by `Stacks population` plugin. We used the `OneMap` wrapper function mds_onemap to order the markers with MDS (Preedy and Hackett, 2016). The genetic distances were estimated by HMM multipoint approach. Figure 7 shows the effects of including the multiallelic markers in the two-points-based MDS algorithm.

**Figure 7:**
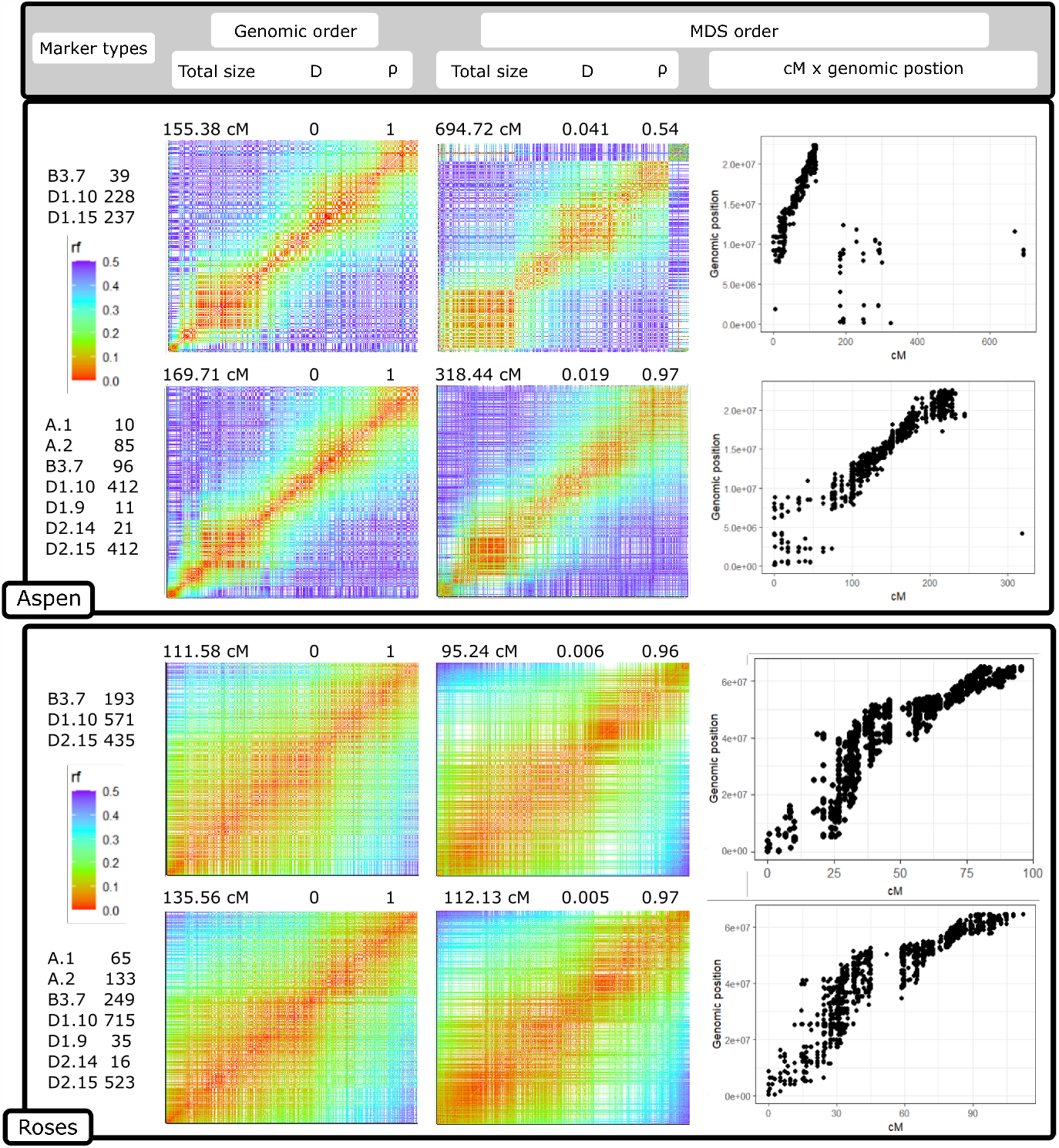
Comparison between MDS ordering algorithm performance in the aspen and rose dataset entire linkage group 10 and 1, respectively with only biallelic markers, and with biallelic and haplotype-based multiallelic markers estimated by `Stacks`. The heatmaps represent the recombination fraction (rf) matrix between markers positioned at both axes. In well-ordered linkage groups, we expect a gradient from hot colors in the diagonal (adjacent markers) to cold colors in the upper left and lower right corners. The figure also presents the Spearman rank correlation (*ρ*) and the Euclidean distances (D) between the estimated map using MDS and the map built with markers ordered by the genomic positions (used as reference). The dot plots relate the positions of markers estimated by MDS with the genomic position.

The impact of multiallelic markers differed between the aspen and roses datasets. In the aspen dataset, characterized by a lower depth and a higher rate of genotyping errors in the markers, most of the B3.7 biallelic markers were filtered out during previous steps, resulting in an unsatisfactory performance of the MDS algorithm in ordering the markers. However, incorporating the multiallelic markers, although slightly inflating the genetic distances, significantly improved the ordering accuracy using MDS. It should be noted that MDS itself can contribute to genetic distance inflation as it may erroneously invert markers in close proximity. In scenarios where a reference genome is unavailable, the inclusion of multiallelic markers can prove valuable for effective marker ordering in these types of datasets.

The rose dataset is characterized by higher-quality markers, and the genomic ordering can be almost entirely reproduced using only biallelic markers. In this scenario, the inclusion of multiallelic markers also leads to a slight inflation of the map size while improving the ordering accuracy through MDS. Unlike the aspen dataset, the MDS algorithm in the rose dataset tends to reduce the genetic distances, resulting in an underestimation of recombination breakpoints. However, considering that there are no significant inversions or translocations (see dot plots in figure 7), we can have more confidence in the genomic order, even if the map is larger. Any discrepancies between the MDS-based order and the genomic order are likely attributed to local changes, which are likely to be errors introduced by MDS.

## 4 Final considerations

The `Reads2Map` workflows have a robust structure to generate production-level results with simple inputs and optimized usage of computational resources. The structure allowed us to test the quality of genetic maps built with the following scenarios: i) using different SNP calling software (`GATK`, `TASSEL`, `Stacks`, and `Freebayes`); ii) using different genotype calling software (`GATK`, `Freebayes`, `TASSEL`, `Stacks`, `updog`, `polyRAD`, `SuperMASSA`); iii) using different linkage map building software (`OneMap` 3.0 and `GUSMap`); iv) establishing different error probabilities (relative to genotype call software, 10%, 1%, 5%, and 0.001% global error, and the combination of the global error rate with the genotype call probabilities); v) applying different marker filtering; vi) with or without multiallelic markers; vi) in empirical and simulated data; vii) with and without segregation distortion; viii) with and without contaminant samples; ix) with different GBS library preparation aspects; and x) with different sequencing depths. These scenarios are commonly found by researchers trying to produce high-quality linkage maps using sequencing technologies. The `Reads2Map` and `Reads2MapApp` are the first tools to guide best practices for building linkage maps with sequencing data pointing software, parameters, and marker filters to be used in diverse scenarios.

We elaborated and limited the scenarios explored according to our experiences as developers of `OneMap`. `OneMap` first version was released in 2007, and since then it has been used to build linkage maps in a diversity of species. Its strategies and structure also served as a base for more complex software such as `MAPpoly` (Mollinari and Garcia, 2019) for building linkage maps in polyploid species. With time, new methods for genetic marker identification using sequencing data emerged, changing the context where `OneMap` was used. We included updates in this version 3.0 to resolve issues with inflated genetic maps and marker ordering. Two major changes allow users to read and build genetic maps with the genotype probabilities and haplotype-based multiallelic markers information from the input files (`OneMap` format or VCF file). However, the success of genetic map building will be proportional to the quality of the information provided by upstream procedures such as library preparation, SNP and genotype calling, genotype probabilities estimation, and the combination of SNPs into haplotype-based markers. With `Reads2Map` and `Reads2MapApp`, we provide users tools to select the best approaches before using `OneMap` 3.0 to guarantee that it will result in the best quality genetic map possible with the data available.

It is important to highlight that we did not design the workflows to be a tool to build a final linkage map but to select the bioinformatic pipeline that provides the best quality genetic markers. Once the pipeline is selected, the respective VCF file and `OneMap` functions can be used in the R environment to build the final map. Building the complete linkage map will require evaluations and edits that are highly specific and cannot be fully automated within the workflows. These tasks include addressing the presence of translocations and inversions, identifying outlier markers, and linkage between markers located in different chromosomes.

The diversity in the results of the pipeline suggested for both empirical datasets highlights that pipelines perform differently with datasets with different properties. This means that the pipelines presented here as the best cannot be considered the best for every dataset. We could reduce the number of required tests by users identifying the dataset-independent approaches and setting them as default in `Reads2Map`. However, we suggest users reproduce the tests presented here for the dataset-dependent approaches using the `Reads2Map` workflows with their empirical dataset and select the best pipelines for their specific conditions.

The workflows were built using WDL and containers to ensure high reproducibility. This guarantees that different results running different datasets is due to the dataset’s properties and not to bioinformatic pipeline changes. Also, updates can be easily made in the workflows as the software implemented are improved once the versions are controlled by Docker images. This makes `Reads2Map` also a useful tool for software developers to validate updates because it facilitates checking the consequences of the changes in the quality of the markers by easily controlling versions, rerunning datasets, and checking the map quality.

Every `Reads2Map` workflow run returns a large amount of information. Every step of the workflow, from the reads’ alignment to the completed linkage map, provides quality measurements for users to evaluate each scenario. The `Reads2MapApp` shiny app receives all this information compressed in a single workflow output file and converts it into comprehensive interactive graphics. Through the app interface, users can evaluate the performance of each combination of software and parameters in each step. If results show issues in any of them, users can re-run the workflow with adapted parameters or include new filters that make sense in their context. Once established the upstream steps based on the app graphics for the built linkage map subset, users can reproduce it for the complete dataset, inputting the VCF files from `Reads2Map` into `OneMap`.

## 5 Availability of source code and requirements

- Project name: Reads2Map
- Project home page: (Taniguti et al., 2022a)
- Main workflows: EmpiricalReads2Map (Taniguti et al., 2022b) and SimulatedReads2Map (Taniguti et al., 2022c)
- Operating system(s): Platform independent
- Programming language: WDL
- Other requirements: docker or singularity
- License: MIT
- RRID: SCR_023593
- biotoolsID: reads2map

## 7 Abbreviations

GBS: Genotyping-by-Sequencing
PCR: polymerase chain reaction
RADSeq: Restrictionsite associated DNA sequencing
VCF: variant call format
GQ: genotyping quality
GT: genotype
GWAS: genome-wide association
SNP: single nucleotide polymorphism
LD: linkage disequilibrium
QTL: quantitative trait loci
WDL: workflow description language
HPRC: high-performance research computing
CPU: central processing unit
HMM: hidden Markov model
EM: expectation-maximization
MAF: minor allele frequency
NGS: Next Generation Sequencing

## 10 Author’s Contributions

CHT, MM, RRA, AAFG, GSP, and GCF contributed to `OneMap` package updates. CHT, LMT, GSG, GSP, AAGF, MM, ORL, and JL contributed with ideas to design `Reads2Map`. CHT and LMT developed and optimized the `Reads2Map` code. CHT developed `Reads2MapApp`. CHT, TPO, AAFG, ORL, DB, and RRA contributed to elaborate the tested scenarios. CHT, TPO, and RRA contributed to analyzing the results. CHT wrote the first version of the manuscript. All authors provided helpful discussions for the work and reviewed the manuscript.

## 11 Acknowledgements

Portions of this research were conducted with the advanced computing resources provided by Texas A&M High-Performance Research Computing and by the University of São Paulo Aguia High-Performance Computing. We also thank David Gerard for the idea of using genotype probabilities from `updog` combined with a global error rate.

## References

Glaubitz JC, Casstevens TM, Lu F, Harriman J, Elshire RJ, Sun Q, et al. TASSEL-GBS: a high capacity genotyping by sequencing analysis pipeline. PLoS ONE 2014 2;9:1–11.

Catchen J, Hohenlohe PA, Bassham S, Amores A, Cresko WA. Stacks: an analysis tool set for population genomics. Molecular Ecology 2013;22:3124–40.

Anderson CB, Franzmayr BK, Hong SW, Larking AC, Stijn TC, Tan R, et al. Protocol: A versatile, inexpensive, high-throughput plant genomic DNA extraction method suitable for genotyping-by-sequencing. Plant Methods 2018 8;14.

Andrews KR, Good JM, Miller MR, Luikart G, Hohenlohe PA. Harnessing the power of RADseq for ecological and evolutionary genomics. Nature Reviews Genetics 2016 1;17:81–92.

Bresadola L, Link V, Buerkle CA, Lexer C, Wegmann D. Estimating and accounting for genotyping errors in RAD-seq experiments. Molecular Ecology Resources 2020;20:856–870.

Baird NA, Etter PD, Atwood TS, Currey MC, Shiver AL, Lewis ZA, et al. Rapid SNP Discovery and Genetic Mapping Using Sequenced RAD Markers. PLoS ONE 2008;3:e3376.

Elshire RJ, Glaubitz JC, Sun Q, Poland JA, Kawamoto K, Buckler ES, et al. A robust, simple genotyping-by-sequencing (GBS) approach for high diversity species. PLoS ONE 2011 5;6:e19379.

der Auwera GV, O’Connor B. Genomics in the Cloud: Using Docker, GATK, and WDL in Terra. O’Reilly Media, Incorporated; 2020.

Rivera-Colón AG, Rochette NC, Catchen JM. Simulation with RADinitio improves RAD-seq experimental design and sheds light on sources of missing data. Molecular Ecology Resources 2020;p. 1–16.

Gerard D, Ferrão LFV, Garcia AAF, Stephens M. Genotyping Polyploids from Messy Sequencing Data. Genetics 2018 11;210:789–807.

a Hackett C, Broadfoot LB. Effects of genotyping errors, missing values and segregation distortion in molecular marker data on the construction of linkage maps. Heredity 2003 1;90:33–38.

Sturtevant AH. The behavior of the chromosomes as studied through linkage. Zeitschrift für induktive Abstammungs-und Vererbungslehre 1915;13:234–287.

Smith GR, Nambiar M. New Solutions to Old Problems: Molecular Mechanisms of Meiotic Crossover Control. Trends in Genetics 2020;36:337–346.

Bilton TP, Schofield MR, Black MA, Chagné D, Wilcox PL, Dodds KG. Accounting for Errors in Low Coverage High-Throughput Sequencing Data When Constructing Genetic Maps Using Biparental Outcrossed Populations. Genetics 2018 5;209:65–76.

Mollinari M, Garcia AAF. Linkage Analysis and Haplotype Phasing in Experimental Autopolyploid Populations with High Ploidy Level Using Hidden Markov Models. G3: Genes|Genomes|Genetics 2019 10;9:3297–3314.

Liao Y, Voorrips RE, Bourke PM, Tumino G, Arens P, Visser RGF, et al. Using probabilistic genotypes in linkage analysis of polyploids. Theoretical and Applied Genetics 2021 8;134:2443–2457.

Margarido GRA, Souza AP, Garcia AAF. OneMap: software for genetic mapping in outcrossing species. Hereditas 2007 7;144:78–9.

Lander ES, Green P. Construction of multilocus genetic linkage maps in humans. Proc Natl Acad Sci USA 1987;84:2363–2367.

Lorenz AJ, Hamblin MT, Jannink JL. Performance of single nucleotide polymorphisms versus haplotypes for genome-wide association analysis in barley. PLoS ONE 2010;5:1–11.

Gawenda I, Thorwarth P, Günther T, Ordon F, Schmid KJ. Genome-wide association studies in elite varieties of German winter barley using single-marker and haplotype-based methods. Plant Breeding 2015 2;134:28–39.

N’Diaye A, Haile JK, Fowler DB, Ammar K, Pozniak CJ. Effect of Co-segregating Markers on High-Density Genetic Maps and Prediction of Map Expansion Using Machine Learning Algorithms. Frontiers in Plant Science 2017 8;8.

Sehgal D, Dreisigacker S. Haplotypes-based genetic analysis: Benefits and challenges. Vav-ilovskii Zhurnal Genetiki i Selektsii 2019;23:803–808.

Abed A, Belzile F. Comparing Single-SNP, Multi-SNP, and Haplotype-Based Approaches in Association Studies for Major Traits in Barley. The Plant Genome 2019;12:190036.

Liu N, Zhang K, in Genetics Zhao HBTA. Haplotype-Association Analysis. Genetic Dissection of Complex Traits 2008;60:335–405.

Jiang Y, Schmidt RH, Reif JC. Haplotype-Based Genome-Wide Prediction Models Exploit Local Epistatic Interactions Among Markers. G3: Genes|Genomes|Genetics 2018;49:g3.300548.2017.

Garrison E, Marth G. Haplotype-based variant detection from short-read sequencing. ArXiv e-prints 2012;p. 9.

McKenna A, Hanna M, Banks E, Sivachenko A, Cibulskis K, Kernytsky A, et al. The Genome Analysis Toolkit: A MapReduce framework for analyzing next-generation DNA sequencing data. Genome Research 2010 9;20:1297–1303.

Clark LV, Lipka AE, Sacks EJ. polyRAD: Genotype Calling with Uncertainty from Sequencing Data in Polyploids and Diploids. G3: Genes|Genomes|Genetics 2019;9:g3.200913.2018.

Serang O, Mollinari M, Garcia AAF. Efficient exact maximum a posteriori computation for bayesian SNP genotyping in polyploids. PLoS ONE 2012;7:1–13.

Patterson M, Marschall T, Pisanti N, van Iersel L, Stougie L, Klau GW, et al. What-sHap: Weighted Haplotype Assembly for Future-Generation Sequencing Reads. Journal of Computational Biology 2015;22:498–509.

Voss K, Gentry J, Auwera GVD. Full-stack genomics pipelining with GATK4+ WDL+ Cromwell [version 1; not peer reviewed]. F1000Research 2017;p. 4.

Taniguti CH, Taniguti LM, Amadeu RR, Lau J, de S Gesteira G, de P Oliveira T, et al. Reads2Map GitHub page 2022;https://github.com/Cristianetaniguti/Reads2Map.

Taniguti CH, Taniguti LM, Amadeu RR, Lau J, de S Gesteira G, de P Oliveira T, et al. EmpiricalReads2Map. WorkflowHub 2022;10.48546/WORKFLOWHUB.WORKFLOW.409.1.

Taniguti CH, Taniguti LM, Amadeu RR, Lau J, de S Gesteira G, de P Oliveira T, et al. SimulatedReads2Map. WorkflowHub 2022;10.48546/WORKFLOWHUB.WORKFLOW.410.1.

bio T. Terra: Focus on your science. Available online at: https://appterrabio/ 2020;.

Merkel D. Docker : Lightweight Linux Containers for Consistent Development and Deployment Docker : a Little Background Under the Hood. Linux Journal 2014;2014:2–7.

Kurtzer GM, Sochat V, Bauer MW. Singularity: Scientific containers for mobility of compute. PLOS ONE 2017 5;12:e0177459.

Taniguti CH, Taniguti LM, Amadeu RR, Lau J, de S Gesteira G, de P Oliveira T, et al. Reads2MapTools GitHub page 2022;https://github.com/Cristianetaniguti/Reads2MapTools.

Taniguti CH, Taniguti LM, Amadeu RR, Lau J, de S Gesteira G, de P Oliveira T, et al. Reads2MapApp GitHub page 2022;https://github.com/Cristianetaniguti/ Reads2MapApp.

Li H. Aligning sequence reads, clone sequences and assembly contigs with BWA-MEM. ArXiv 2013;1303.

Li H, Handsaker B, Wysoker A, Fennell T, Ruan J, Homer N, et al. The sequence alignment/map format and SAMtools. Bioinformatics 2009;25:2078–2079.

Danecek P, Bonfield JK, Liddle J, Marshall J, Ohan V, Pollard MO, et al. Twelve years of SAMtools and BCFtools. GigaScience 2021 1;10.

Knaus BJ, Grünwald NJ. vcfR: a package to manipulate and visualize variant call format data in R. Molecular Ecology Resources 2017 1;17:44–53.

Baum E, Petrie T G S N W. A Maximization Technique Occurring in the Statistical Analysis of Probabilistic Functions of Markov Chains. The Annals of Mathematical Statistics 1970;41:164–171.

Schiffthaler B, Bernhardsson C, Ingvarsson PK, Street NR. BatchMap: A parallel implementation of the OneMap R package for fast computation of F1 linkage maps in outcrossing species. PLoS ONE 2017;12:1–12.

Guyader V, Fay C, Rochette S, Girard C. golem: A Framework for Robust Shiny Applications. Golem GitHub repository 2022;https://github.com/ThinkR-open/golem.

Zhigunov AV, Ulianich PS, Lebedeva MV, Chang PL, Nuzhdin SV, Potokina EK. Development of F1 hybrid population and the high-density linkage map for European aspen (Populus tremula L.) using RADseq technology. BMC Plant Biology 2017;17.

Young EL, Lau J, Bentley NB, Rawandoozi Z, Collins S, Windham MT, et al. Identification of QTLs for Reduced Susceptibility to Rose Rosette Disease in Diploid Roses. Pathogens 2022 6;11:660.

Tuskan GA, DiFazio S, Jansson S, Bohlmann J, Grigoriev I, Hellsten U, et al. The Genome of Black Cottonwood, Populus trichocarpa. Science 2006 9;313:1596–1604.

Saint-Oyant LH, Ruttink T, Hamama L, Kirov I, Lakhwani D, Zhou NN, et al. A high-quality genome sequence of Rosa chinensis to elucidate ornamental traits. Nature Plants 2018 7;4:473–484.

Martin M. Cutadapt removes adapter sequences from high-throughput sequencing reads. EMBnetjournal 2011 5;17:10.

Hyman JM. Accurate Monotonicity Preserving Cubic Interpolation. SIAM Journal on Scientific and Statistical Computing 1983 12;4:645–654.

Wu R, Ma CX, Wu SS, Zeng ZB. Linkage mapping of sex-specific differences. Genetical research 2002;79:85–96.

Voorrips RE, Maliepaard CA. The simulation of meiosis in diploid and tetraploid organisms using various genetic models. BMC Bioinformatics 2012 12;13:248.

Haldane JBS. The combination of linkage values, and the calculation of distance between linked factors. Journal of Genetics 1919;8:299–309.

Best K, Oakes T, Heather JM, Shawe-Taylor J, Chain B. Computational analysis of stochastic heterogeneity in PCR amplification efficiency revealed by single molecule barcoding. Scientific reports 2015 10;5:14629.

Glenn TC. Field guide to next-generation DNA sequencers. Molecular Ecology Resources 2011;11:759–769.

Li H. seqtk: Toolkit for processing sequences in FASTA/Q formats. seqtk GitHub repository 2020;https://github.com/lh3/seqtk.

Ros-Freixedes R, Gonen S, Gorjanc G, Hickey JM. A method for allocating low-coverage sequencing resources by targeting haplotypes rather than individuals. Genetics Selection Evolution 2017;49:1–17.

Preedy KF, Hackett CA. A rapid marker ordering approach for high-density genetic linkage maps in experimental autotetraploid populations using multidimensional scaling. Theoretical and Applied Genetics 2016;.

DePristo MA, Banks E, Poplin R, Garimella KV, Maguire JR, Hartl C, et al. A framework for variation discovery and genotyping using next-generation DNA sequencing data. Nature Genetics 2011;43:491–8.

Duncavage EJ, Coleman JF, de Baca ME, Kadri S, Leon A, Routbort M, et al. Recommendations for the Use of In silico Approaches for Next Generation Sequencing Bioinformatic Pipeline Validation: A Joint Report of the Association for Molecular Pathology, Association for Pathology Informatics, and College of American Pathologists. The Journal of molecular diagnostics : JMD 2022 10;.

Alexander DH, Novembre J, Lange K. Fast model-based estimation of ancestry in unrelated individuals. Genome Research 2009 9;19:1655–1664.

Amadeu RR, Cellon C, Olmstead JW, Garcia AAF, Resende MFR, Muñoz PR. AGHmatrix: R Package to Construct Relationship Matrices for Autotetraploid and Diploid Species: A Blueberry Example. The Plant Genome 2016 11;9.

Grattapaglia D, Sederoff R. Genetic linkage maps of Eucalyptus grandis and Eucalyptus urophylla using a pseudo-testcross: mapping strategy and RAPD markers. Genetics 1994 8;137:1121–37.

Gazaffi R, Margarido GRA, Pastina MM, Mollinari M, Garcia AAF. A model for quantitative trait loci mapping, linkage phase, and segregation pattern estimation for a full-sib progeny. Tree Genetics and Genomes 2014;10:791–801.

