## Supplementary material for "Developing best practices for genotyping-by-sequencing analysis in the construction of linkage maps": Figures and tables

#### Supplementary File 1 - Genotype probabilities in OneMap 3.0 Hidden Markov Model

With a combination of a hidden Markov model (HMM) and the expectation-maximization algorithm (EM) (Lander and Green, 1987), OneMap (Margarido et al., 2007) can perform multipoint estimation of map genetic distance for F2, backcross, RILs, and outcrossing populations. For the multipoint estimation, OneMap algorithms use code adapted from R/QTL package (Broman et al., 2003).

In short, the latent variable $G_i, i = 1, ..., n$, denotes the true underlying genotypes for the individual at a set of $n$ ordered loci; $O_i$ is the observed variable of the molecular phenotype (observed genotypes) for the locus $i$. The HMM can be represented as (Broman et al., 2009):

$$P(O|G_i = g_i) = \sum_{g_1} ... \sum_{g_i - 1} \sum_{g_i + 1} ... \sum_{g_n} \pi(g_1) \prod_{j=1}^{n-1} t_j(g_j, g_{j+1}) \prod_{j=1}^{n} e(g_j, O_j) \quad (1)$$

The initial probability $\pi(g_1)$ is the probability of having a given genotype for the first locus ($G_1$), and its value depends on the cross-type. For example, for an outcrossing population, this value will be 0.25, assuming a uniform distribution of all four possible genotypes (AA, BA, AB, and BB). The same reasoning applies to backcross data, with probabilities of 0.5 since there are only two possible genotypes (AA and AB).

The transition probability $t_j(g_j, g_{j+1})$ is the probability of the genotype in a locus ($G_{j=i+1}$) changing to the next locus genotype ($G_{j+1}$). The initial value for this probability is based on the phase, and recombination fraction estimated by a two-point approach using maximum likelihood estimators (Maliepaard et al., 1997), and is updated after iterations of the EM algorithm.

The emission probability $e(g_j, O_j)$ is the probability of the observed variable given the genotype, it considers a probability for every possible phased genotype and can include an associated genotyping error. For outcrossing and $F_2$ intercrossing, there are four possibilities. Here we will denote them generically as "AA", "AB", "BA", and "BB". The probability value that each one will receive depends on which genotype was observed, the marker type, and the associated error of the genotype ($e$). As an example, If we have a marker type A ("ab" x "cd") we can observe four different genotypes, if we observe the genotype "ac" with high confidence ($e = 0$), the phase "AA" would receive the maximum probability (1) and the others would receive 0. In the older version, OneMap considered a unique error of $10^{-5}$ ($e$), which means that if the genotype was called as "ac", the "AA" phased genotype has a probability of $1 - 10^{-5}$ and the others have $10^{-5}/3$. This way, we can include an uncertainty between the observed genotype and the estimated phased genotype, which characterizes the hidden aspect of the HMM.

For marker type A, this could not seem very useful because the genotype phase is already represented in the observed genotypes. But other marker types do not have a direct relationship between the observed genotype and the estimated phased genotype. For example, if we have a marker type B3.7 and observe the "ab" genotype, the estimated phased genotype can be the "AB" or "BA", and we will consider equal probabilities for them in the emission function. The multipoint aspect of the HMM combined with the expectation-maximization

(EM) will change these probabilities, and, in the end, we will be able to differentiate between phased genotypes.

In `OneMap` 3.0, users can now provide customized error rates or genotype probabilities to control specific errors in their dataset. The values defined by users will be applied in three different ways in the emission function of the HMM. The variable $e$ represents the error rate described in equation 3.1. If users define a single value (*global_error* argument), it will be the error rate for all observed genotypes. If users provide the genotype errors (*genotype_errors*), each genotype observed can receive a different error rate value. If users provide a probability for each genotype (*genotypes_probs*), the values in each cell of the following table will be replaced by their respective user-provided genotype probability. The tables below are based on R/QTL (Broman et al., 2003) emission function and describe how the error values are implemented in `OneMap` HMM.

Each observed genotype (columns) receives specific genotype probabilities according to marker type segregation and the error rate. The emission function returns from the iterative steps of the HMM the estimated probability for the phased genotypes. In general, the error rate allows the HMM to change the genotypes according to the information from the entire sequence (Mollinari and Garcia, 2019) or batches with proper size (Schiffthaler et al., 2017).

Table S1: Emission function values according to marker types. The error rate is represented by $e$, unphased genotypes as "a", "b", "c", "d" and their combination. The estimated phased genotype s are represented by "AA", "AB", "BA" and "BB". The "o" represents null alleles. Marker types follow the segregation pattern as described in Wu et al. (2002)

| Marker type | A | observed genotypes | | | |
|---|---|---|---|---|---|
| Marker sub-types | A1 | ac | ad | bc | bd |
| | A2 | a | ac | ba | bc |
| | A3 | ac | a | bc | b |
| | A4 | ab | a | b | o |
| Estimated phased genotypes | AA | 1-$e$ | $e/3$ | $e/3$ | $e/3$ |
| | AB | $e/3$ | 1-$e$ | $e/3$ | $e/3$ |
| | BA | $e/3$ | $e/3$ | 1-$e$ | $e/3$ |
| | BB | $e/3$ | $e/3$ | $e/3$ | 1-$e$ |
| OneMap codification | | 1 | 2 | 3 | 4 |

Table S2: Continued from table

| Marker type | B1 | observed genotypes | | |
|---|---|---|---|---|
| Marker sub-types | B1.5 | a | ab | b |
| Estimated phased genotypes | AA | 1-$e$ | $e/3$ | $e/3$ |
| | AB | 1-$e$ | $e/3$ | $e/3$ |
| | BA | $e$ | 1-$e$ | $e/3$ |
| | BB | $e$ | $e/3$ | 1-$e$ |
| OneMap codification | | 1 | 2 | 3 |

Table S5: Continued from table

| Marker type | C | observed genotypes | |
|---|---|---|---|
| Marker sub-types | C.8 | a | o |
| Estimated phased genotypes | AA | (1-$e$)/3 | $e$/3 |
| | AB | (1-$e$)/3 | $e$/3 |
| | BA | (1-$e$)/3 | $e$/3 |
| | BB | $e$ | 1-$e$ |
| OneMap codification | | 1 | 2 |

Table S3: Continued from table

| Marker type | B2 | observed genotypes | | |
|---|---|---|---|---|
| Marker sub-types | B2.6 | a | ab | b |
| Estimated phased genotypes | AA | 1-$e$ | $e$/3 | $e$/3 |
| | AB | $e$ | 1-$e$ | $e$/3 |
| | BA | 1-$e$ | $e$/3 | $e$/3 |
| | BB | $e$ | $e$/3 | 1-$e$ |
| OneMap codification | | 1 | 2 | 3 |

Table S4: Continued from table

| Marker type | B3 | observed genotypes | | |
|---|---|---|---|---|
| Marker sub-types | B3.7 | a | ab | b |
| Estimated phased genotypes | AA | 1-$e$ | $e$ | $e$/3 |
| | AB | $e$/3 | 1-$e$ | $e$/3 |
| | BA | $e$/3 | 1-$e$ | $e$/3 |
| | BB | $e$/3 | $e$ | 1-$e$ |
| OneMap codification | | 1 | 2 | 3 |

Table S6: Continued from table

| Marker type | D1 | observed genotypes | |
|---|---|---|---|
| Marker sub-types | D1.9 | ac | bc |
| | D1.10 | a | ab |
| | D1.11 | a | b |
| | D1.12 | ab | a |
| | D1.13 | a | o |
| Estimated phased genotypes | AA | 1-$e$ | $e$ |
| | AB | 1-$e$ | $e$ |
| | BA | $e$ | 1-$e$ |
| | BB | $e$ | 1-$e$ |
| OneMap codification | | 1 | 2 |

Table S7: Continued from table

| Marker type | D2 | observed genotypes | |
|---|---|---|---|
| Marker sub-types | D2.14 | ac | bc |
| | D2.15 | a | ab |
| | D2.16 | a | b |
| | D2.17 | ab | a |
| | D2.18 | a | o |
| Estimated phased genotypes | AA | 1-$e$ | $e$ |
| | AB | $e$ | 1-$e$ |
| | BA | 1-$e$ | $e$ |
| | BB | $e$ | 1-$e$ |
| OneMap codification | | 1 | 2 |

Supplementary File 2 - `Reads2MapApp` interface demonstration

To access `Reads2MapApp`, install and run it in the R environment:

```
library(devtools)
install_github("Cristianetaniguti/Reads2MapApp")
Reads2MapApp::run_app()
```

The command will make the app interface pop up. The first page has a short description of the app's functionalities. Other page options can be accessed by clicking on the menu icon.

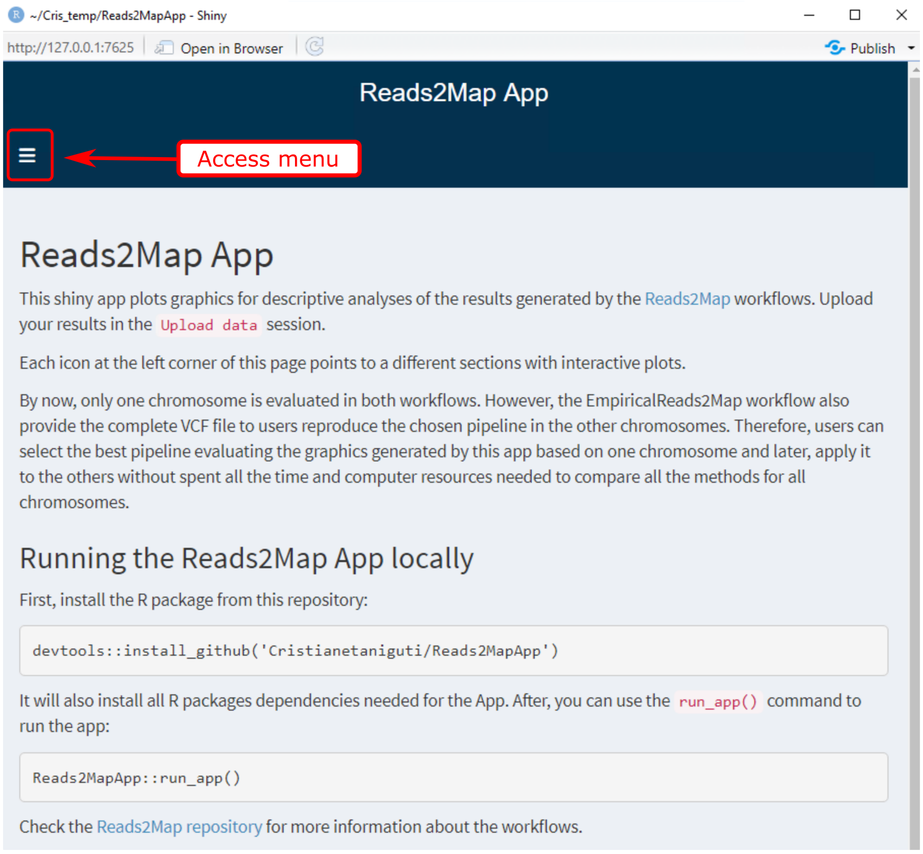


Figure S1: `Reads2MapApp` about page. The red arrow indicates the menu icon to access the app's other pages.

Users can upload their data in the "Upload data" section:

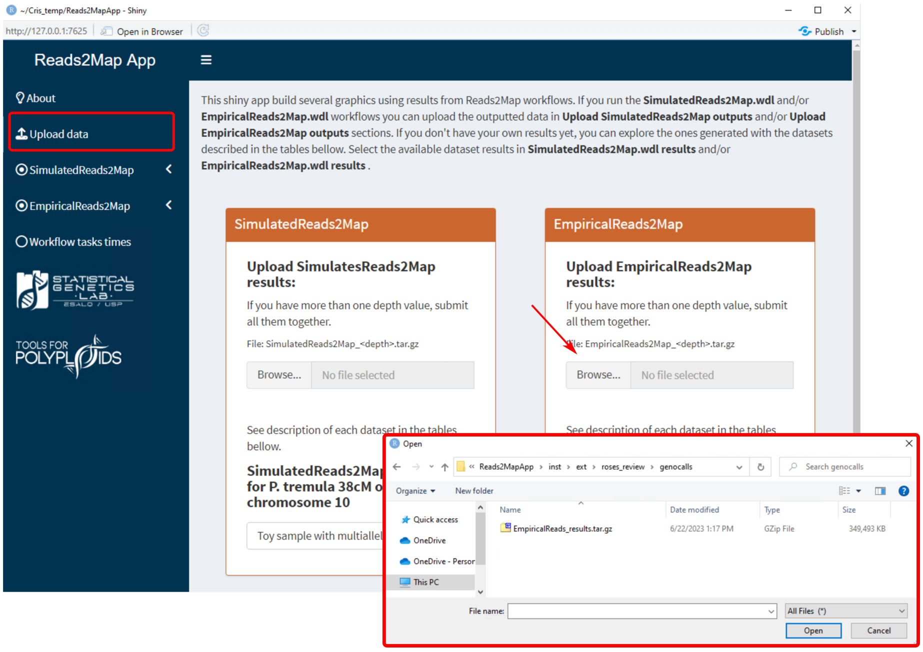

Figure S2: `Reads2MapApp` upload page. The red arrow indicates the button to upload the EmpiricalMaps workflow results.

Once uploaded the options available in each one of the "Empiricalreads2Map" sections will be updated according to the results of the workflow.

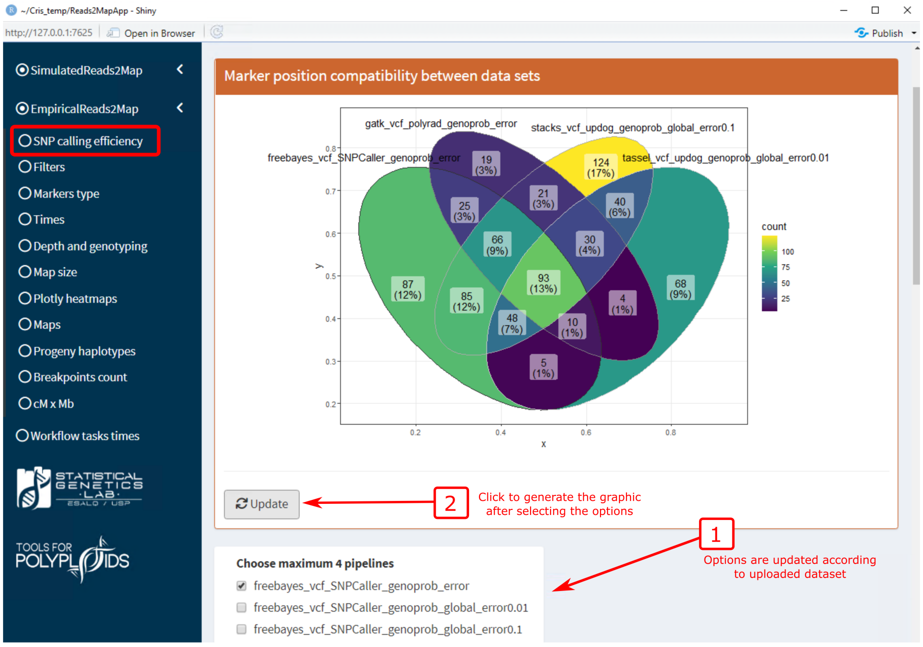


Figure S3: Example of the "SNP calling efficiency" section. Venn diagrams are built to show the number of markers identified in the pipelines defined in the options and the common markers between them.

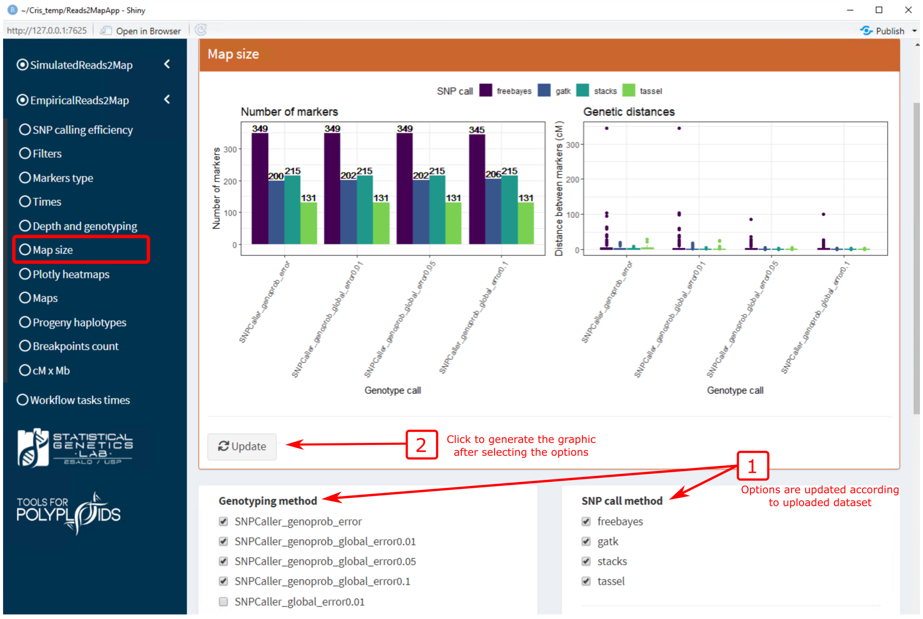


Figure S4: Example of the "Map size" section of `Reads2MapApp`. The graphic shows the number of markers (left) and the distances between adjacent markers (right) for each method.

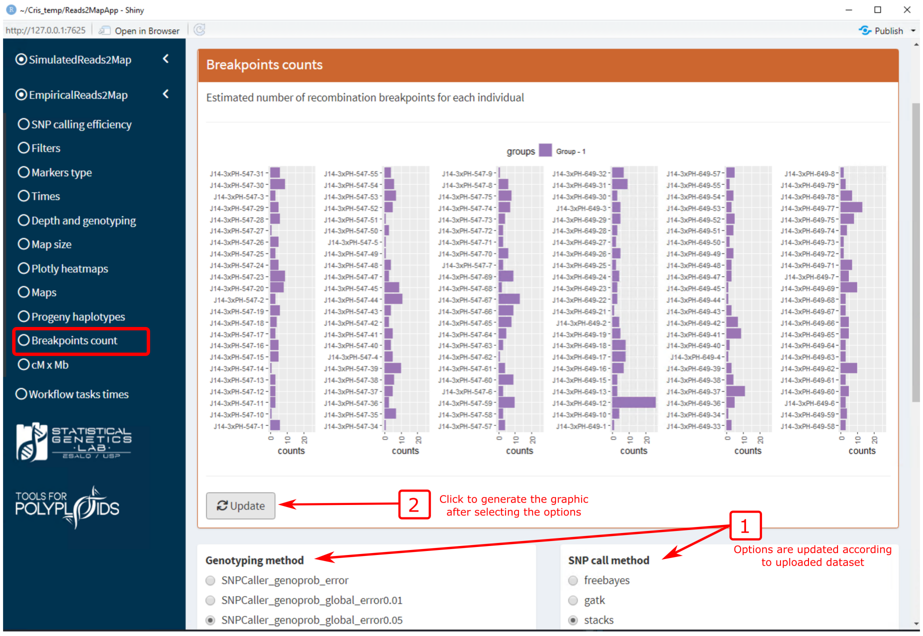


Figure S5: Example of the "Breakpoint count" section of `Reads2MapApp`. The graphic shows the number of estimated breakpoints in each progeny haplotype according to the method selected in the options.

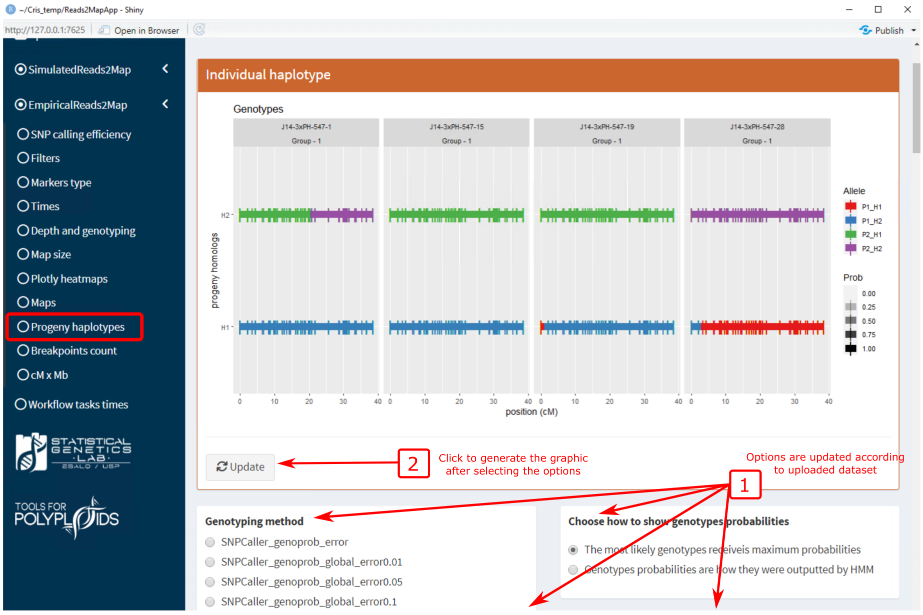


Figure S6: Example of the "Progeny haplotypes" section of `Reads2MapApp`. The graphic shows the estimated haplotype for the individuals selected in the "Individuals from progeny" options according to the method selected in the options.

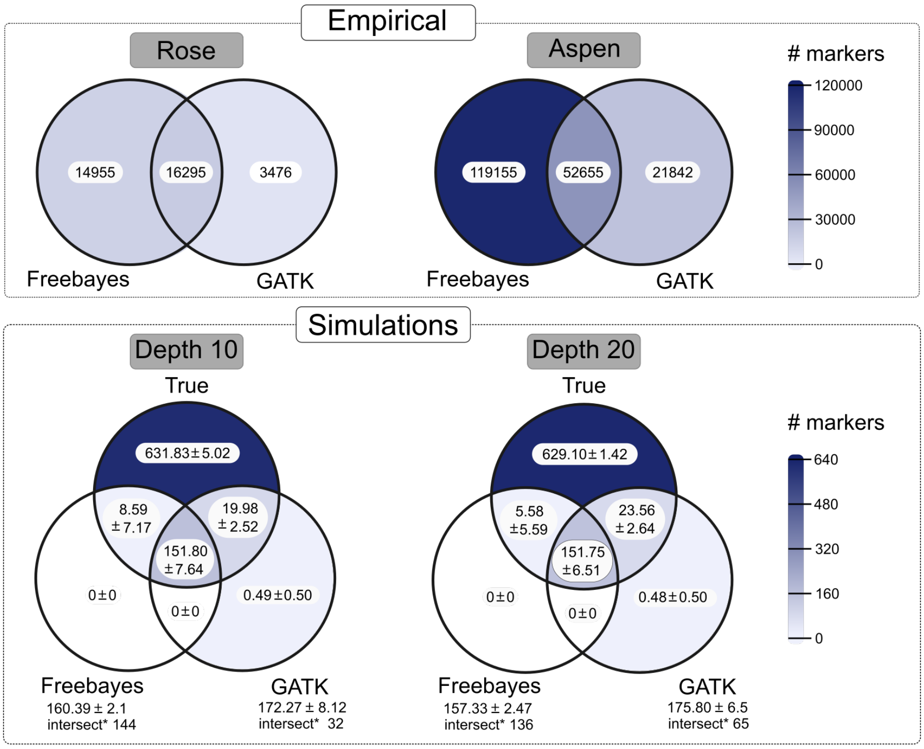


Figure S7: Venn diagrams show the number of markers identified by `freebayes`, `GATK`, and simulated (true). The intersection between the data sets represents markers with the same position in the reference genome *Populus trichocarpa* version 3.0. The Empirical data sets include markers spread across the entire reference genome. The simulations only include markers in the first 8.426 Mb of chromosome 10 (2.1% of the genome). The mean and standard deviation of number markers are shown for the simulated data set once the simulation and SNP calling are repeated 60 times. Markers were filtered by 25% maximum missing data and MAF 5% in empirical and simulated data. * Number of markers common to all 60 repetitions.

Supplementary Table 8 - List of third-party software and images versions used

| Software | Reference | Image |
|---|---|---|
| BWA | Li (2013) | us.gcr.io/broad-gotc-prod/genomes-in-the-cloud:2.5.7-2021-06-09_16-47-48Z |
| cutadapt | Martin (2011) | cristaniguti/pirs-ddrad-cutadapt:0.0.1 |
| Freebayes | Garrison and Marth (2012) | Cristaniguti/freebayes:0.0.1 |
| GATK | McKenna et al. (2010) | us.gcr.io/broad-gotc-prod/genomes-in-the-cloud:2.5.7-2021-06-09_16-47-48Z |
| TASSEL | Glaubitz et al. (2014) | cristaniguti/java-in-the-cloud:0.0.2 |
| STACKs | Catchen et al. (2013) | cristaniguti/stacks:0.0.1 |
| PedigreeSim | Voorrips and Maliepaard (2012) | cristaniguti/reads2map:0.0.1 |
| picard | Institute (2009) | us.gcr.io/broad-gotc-prod/genomes-in-the-cloud:2.5.7-2021-06-09_16-47-48Z |
| samtools | Li et al. (2009) | us.gcr.io/broad-gotc-prod/genomes-in-the-cloud:2.5.7-2021-06-09_16-47-48Z |
| RADinitio | Rivera-Colón et al. (2020) | cristaniguti/radinitio:0.0.1 |
| SuperMASSA | Serang et al. (2012) | cristaniguti/reads2map:0.0.1 |
| bcftools | Danecek et al. (2021) | lifebitai/bcftools:1.10.2 |
| OneMap | Margarido et al. (2007) and updated in this work | cristaniguti/reads2map:0.0.1 |
| Reads2MapTools | Developed in this work | cristaniguti/reads2map:0.0.1 |
| GUSMap | Bilton et al. (2018) | cristaniguti/reads2map:0.0.1 |
| updog | Gerard et al. (2018) | cristaniguti/reads2map:0.0.1 |
| polyRAD | Clark et al. (2019) | cristaniguti/reads2map:0.0.1 |
| Reads2MapApp | Developed in this work | cristaniguti/reads2mapApp:0.0.1 |
| simuscopR | Developed in this work | cristaniguti/reads2map:0.0.1 |

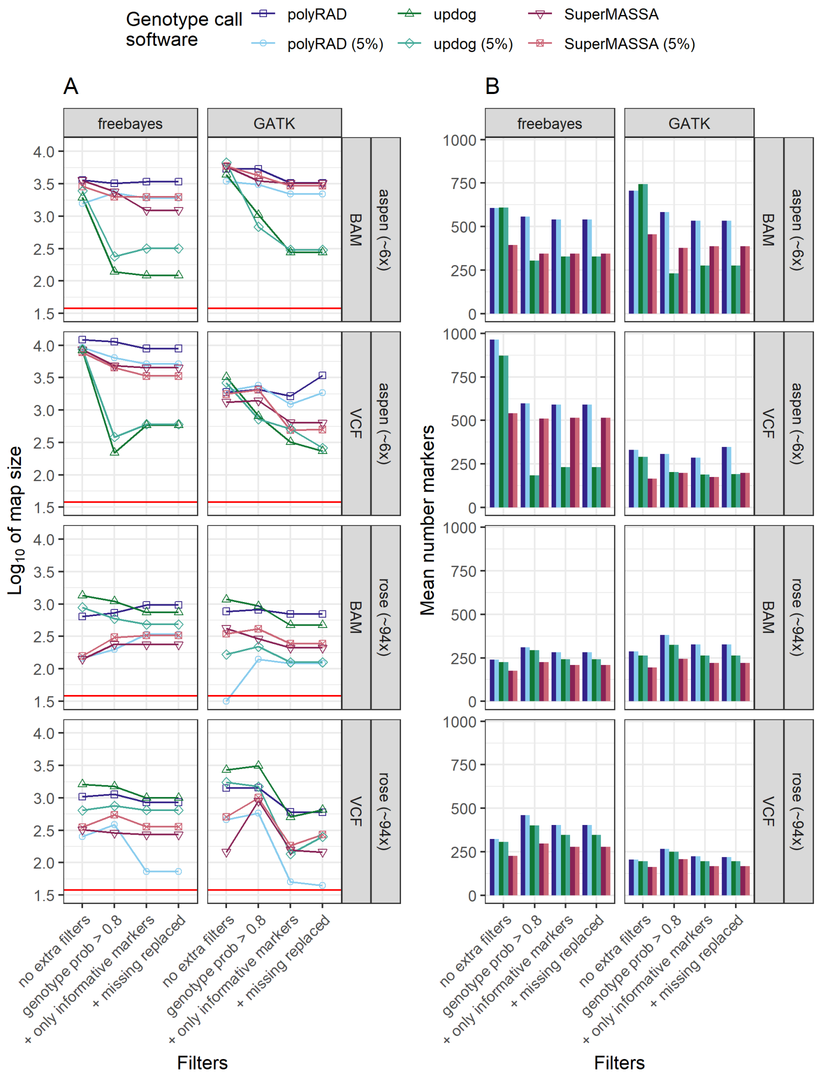


Figure S8: The relation between filters applied (x-axis), the map size (A y-axis), and the number of markers (B y-axis) for genotype calling software used in the empirical data sets. The data sets shown in the figure contain only biallelic markers. The horizontal red line indicates the expected map size (38 cM) for the subset of the genomes used.

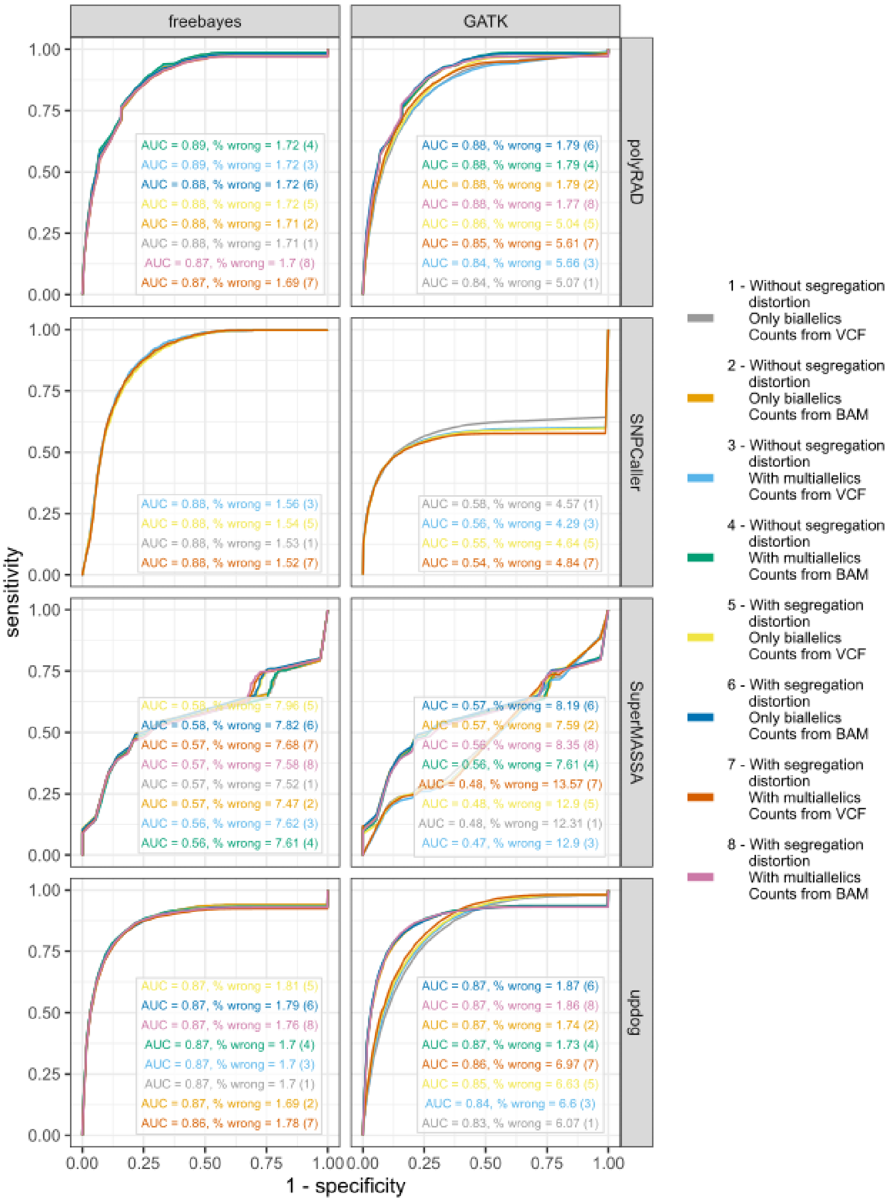


Figure S9: ROC curves with the true and estimated genotypes from the five families simulated with mean depth 10 and 20 and with the firsts 8.426 Mb of the chromosome 10 (37% or 38 cM). Here only biallelic markers are considered. The specificity and sensitivity profiles consider different thresholds in the genotype probabilities for each scenario. The higher the area under the curve, the higher the genotype's probability reliability. Genotype probabilities thresholds closer to the left superior corner have a higher capacity to differentiate right and wrong genotypes.

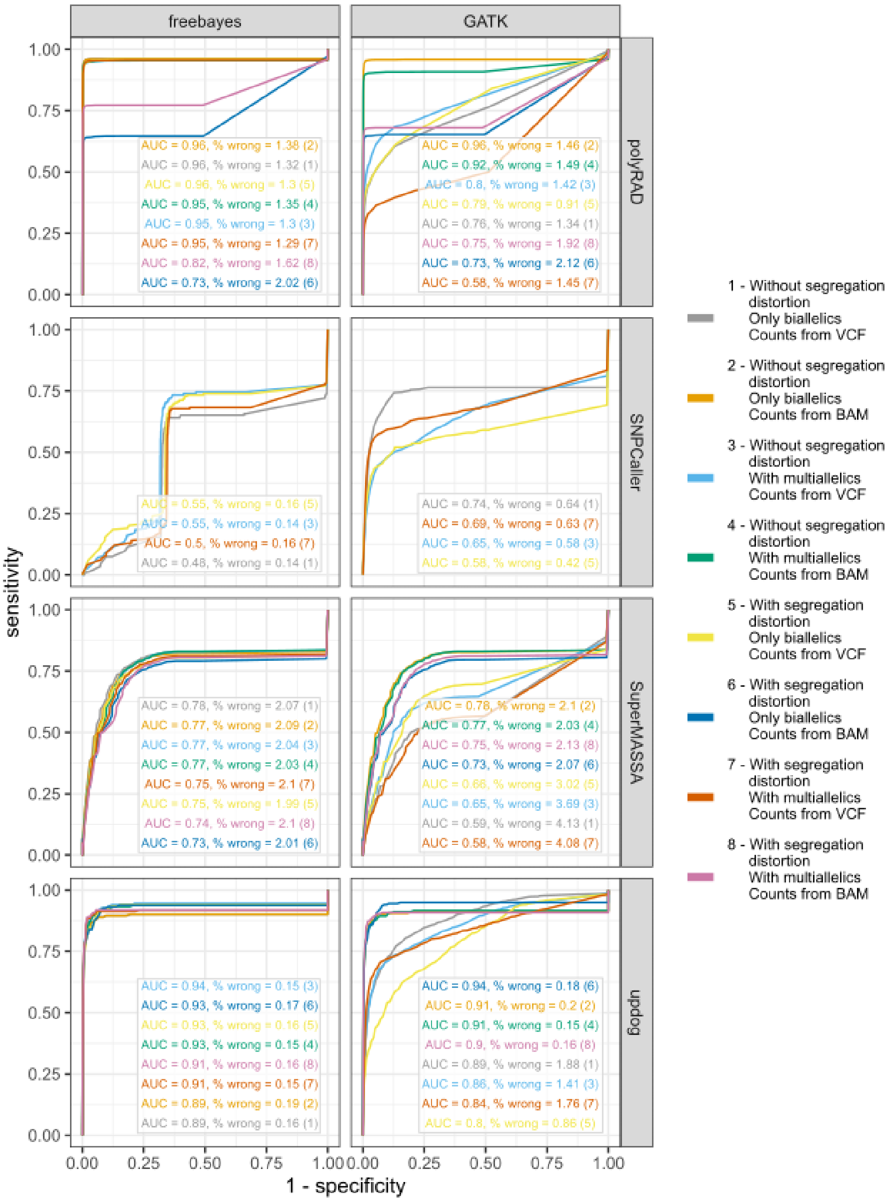


Figure S10: See supplementary figure S9 description.

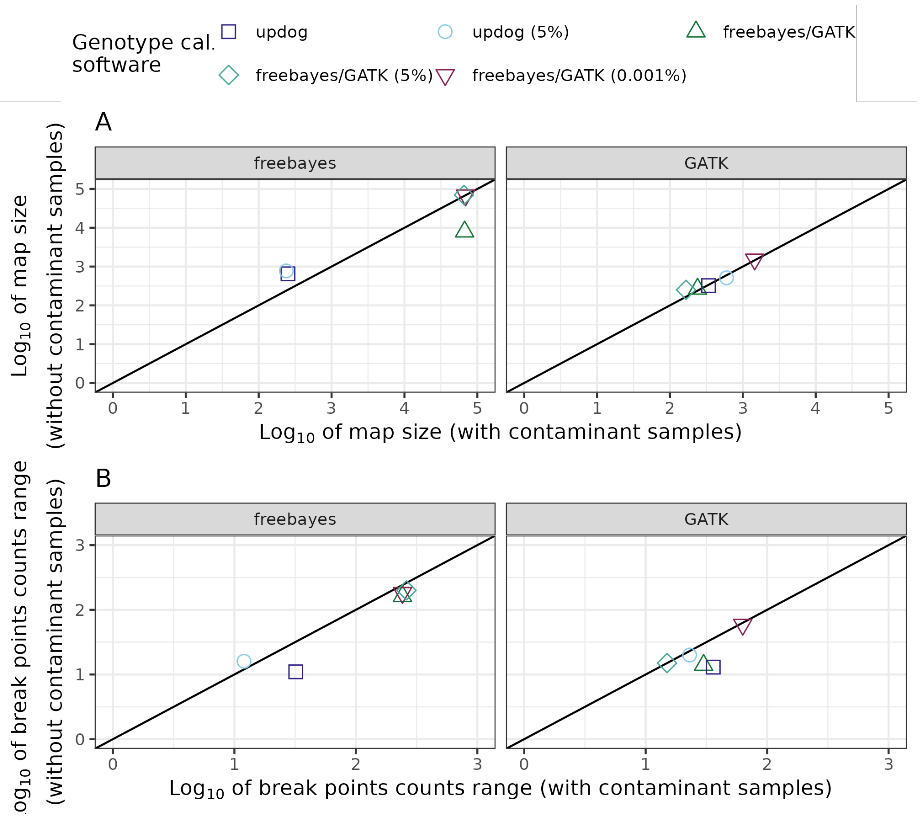


Figure S11: Effect of contaminant samples in the map size (A) and in the number of estimated recombination breakpoints range (B) among progeny individuals. The empirical aspen data sets presented in this figure contain multiallelic markers, the allele counts from the VCF file, and is filtered by genotype probability higher than 0.8 to keep only informative markers.

### References


Lander ES, Green P. Construction of multilocus genetic linkage maps in humans. Proc Natl Acad Sci USA 1987;84:2363–2367.

Margarido GRA, Souza AP, Garcia AAF. OneMap: software for genetic mapping in outcrossing species. Hereditas 2007 7;144:78–9.

Broman KW, Wu H, Sen S, Churchill GA. R/qtl: QTL mapping in experimental crosses. Bioinformatics 2003;19:889–890.

Broman KW, Śaunak Sen, Sen S. A Guide to QTL Mapping with R/qtl, vol. 66. Springer New York; 2009.

Maliepaard C, Jansen J, Ooijen JWV. Linkage analysis in a full-sib family of an outbreeding plant species : overview and consequences for applications. Genetical Research 1997;70:237–250.

Mollinari M, Garcia AAF. Linkage Analysis and Haplotype Phasing in Experimental Autopolyploid Populations with High Ploidy Level Using Hidden Markov Models. G3: Genes—Genomes—Genetics 2019 10;9:3297–3314.

Schiffthaler B, Bernhardsson C, Ingvarsson PK, Street NR. BatchMap: A parallel implementation of the OneMap R package for fast computation of F1 linkage maps in outcrossing species. PLoS ONE 2017;12:1–12.

Wu R, Ma CX, Painter I, Zeng ZB. Simultaneous Maximum Likelihood Estimation of Linkage and Linkage Phases in Outcrossing Species. Theoretical Population Biology 2002 5;61:349–363.

Li H. Aligning sequence reads, clone sequences and assembly contigs with BWA-MEM. ArXiv 2013;1303.

Martin M. Cutadapt removes adapter sequences from high-throughput sequencing reads. EMBnetjournal 2011 5;17:10.

Garrison E, Marth G. Haplotype-based variant detection from short-read sequencing. ArXiv e-prints 2012;p. 9.

McKenna A, Hanna M, Banks E, Sivachenko A, Cibulskis K, Kernytsky A, et al. The Genome Analysis Toolkit: A MapReduce framework for analyzing next-generation DNA sequencing data. Genome Research 2010 9;20:1297–1303.

Glaubitz JC, Casstevens TM, Lu F, Harriman J, Elshire RJ, Sun Q, et al. TASSEL-GBS: a high capacity genotyping by sequencing analysis pipeline. PLoS ONE 2014 2;9:1–11.

Catchen J, Hohenlohe PA, Bassham S, Amores A, Cresko WA. Stacks: an analysis tool set for population genomics. Molecular Ecology 2013;22:3124–40.

Voorrips RE, Maliepaard CA. The simulation of meiosis in diploid and tetraploid organisms using various genetic models. BMC Bioinformatics 2012 12;13:248.

Institute B. Picard Tools. Broad Institute, GitHub repository 2009;https://github.com/broadinstitute/picard.

Li H, Handsaker B, Wysoker A, Fennell T, Ruan J, Homer N, et al. The sequence alignment/map format and SAMtools. Bioinformatics 2009;25:2078–2079.

Rivera-Colón AG, Rochette NC, Catchen JM. Simulation with RADinitio improves RADseq experimental design and sheds light on sources of missing data. Molecular Ecology Resources 2020;p. 1–16.

Serang O, Mollinari M, Garcia AAF. Efficient exact maximum a posteriori computation for bayesian SNP genotyping in polyploids. PLoS ONE 2012;7:1–13.

Danecek P, Bonfield JK, Liddle J, Marshall J, Ohan V, Pollard MO, et al. Twelve years of SAMtools and BCFtools. GigaScience 2021 1;10.

Bilton TP, Schofield MR, Black MA, Chagné D, Wilcox PL, Dodds KG. Accounting for Errors in Low Coverage High-Throughput Sequencing Data When Constructing Genetic Maps Using Biparental Outcrossed Populations. Genetics 2018 5;209:65–76.

Gerard D, Ferrão LFV, Garcia AAF, Stephens M. Genotyping Polyploids from Messy Sequencing Data. Genetics 2018 11;210:789–807.

Clark LV, Lipka AE, Sacks EJ. polyRAD: Genotype Calling with Uncertainty from Sequencing Data in Polyploids and Diploids. G3: Genes—Genomes—Genetics 2019;9:g3.200913.2018.
